# A blueprint for national assessments of the blue carbon capacity of kelp forests applied to Canada’s coastline

**DOI:** 10.1101/2024.04.05.586816

**Authors:** Jennifer McHenry, Daniel K. Okamoto, Karen Filbee-Dexter, Kira Krumhansl, Kathleen A. MacGregor, Margot Hessing-Lewis, Brian Timmer, Philippe Archambault, Claire M. Attridge, Delphine Cottier, Maycira Costa, Matt Csordas, Ladd E. Johnson, Joanne Lessard, Alejandra Mora-Soto, Anna Metaxas, Chris Neufeld, Ondine Pontier, Luba Reshitnyk, Samuel Starko, Jennifer Yakimishyn, Julia K. Baum

## Abstract

Kelp forests offer substantial carbon fixation, with the potential to contribute to natural climate solutions (NCS). However, to be included in national NCS inventories, governments must first quantify the kelp-derived carbon stocks and fluxes leading to carbon sequestration. Here, we present a blueprint for assessing the national carbon sequestration capacity of kelp forests in which data synthesis and Bayesian hierarchical modelling enable estimates of kelp forest carbon production, storage, and export capacity from limited data. Applying this blueprint to Canada’s extensive coastline, we find kelp forests store an estimated 1.4 Tg C in short-term biomass and produce 3.1 Tg C yr^-1^ with modest carbon fluxes to the deep ocean. Arctic kelps had the highest carbon stocks and production capacity, while Pacific kelps had greater carbon fluxes overall due to their higher productivity and export rates. Our transparent, reproducible blueprint represents an important step towards accurate carbon accounting for kelp forests.

## Introduction

As the urgency of addressing climate change intensifies, natural climate solutions (NCS) involving habitat interventions to enhance natural carbon sinks have emerged as distinct components of countries’ mitigation strategies^1,2^. However, most NCS assessments focus on forests, grasslands, and wetlands, with less attention on the vast carbon reservoirs found in the ocean^1,3,4^. In the coastal zone, blue carbon ecosystems (BCEs)–seagrass meadows, salt marshes, and mangrove forests–contribute to carbon sequestration in the ocean by converting dissolved carbon dioxide (CO_2_) that has been removed from the atmosphere into biomass, and by promoting the burial of organic material in benthic sediments^2,5–8^. BCE standing biomass can persist for decades, and sedimentary carbon stocks can be preserved for centuries to millennia when undisturbed^9–11^. As a result, these systems remove carbon from shallow waters where it would have otherwise exchanged as atmospheric CO_2_ and exacerbated climate change^8^. Since many BCEs have declined significantly over the past century^12^, conservation and improved management of these ecosystems are increasingly seen as low regret strategies for avoiding further CO_2_ emissions. Similarly, restoration and expansion of BCEs has also been proposed as a potential strategy to enhance natural carbon sequestration in the ocean^1,2,13^.

Kelp forests, composed of large brown seaweeds primarily from the order Laminariales, have traditionally not been considered blue carbon ecosystems, due to their lack of roots and local carbon burial in sediments^14,15^. However, recent work identifies kelp forests as emerging BCEs^16^ because of their ability to efficiently assimilate CO_2_ ^17^, their near global distributions^18,19^, their role as allochthonous producers of carbon-rich material, and their potential export to depositional environments where sequestration occurs ^20–23^. Much like terrestrial forests, kelps form expansive and highly productive vegetated canopies, with some species extending from the benthos to the surface (i.e., surface kelps) and others forming dense submerged beds on the seafloor (i.e., subsurface kelps). While most kelp production enters marine food webs as particulate and dissolved organic carbon (POC and DOC, respectively) and is remineralized in the short-term ^24^, a portion has the potential to become sequestered and stored for geological timescales (i.e., 100s to 1000s of years) in various natural ocean carbon sinks^14,17,25^. There are three main pathways for kelp carbon sequestration: *1)* some portion of kelp DOC is or becomes refractory DOC (i.e., inaccessible to microbial communities) with residence times ranging from decades to millennia when exported below the photic zone^20,26^; *2)* kelp POC in the form of dislodged or fragmented kelp fronds is transported and buried in shelf sediments and the sediments of other BCEs (e.g., seagrass meadows) for similar timescales ^27,28^; and 3) kelp POC and DOC reaches the deep ocean (depths >200 m), where if buried can be preserved for centuries to millennia because of the limited potential for resuspension to the surface ocean^20,29^.

Global assessments show considerable potential for carbon assimilation through kelp productivity ^17,18^. Yet whether kelp forests can provide viable NCS remains unclear due to the data gaps, process uncertainties, and the challenges associated with measuring kelp carbon sequestration at relevant scales for management (e.g., regional or national)^30^. Substantial stretches of temperate and sub-arctic coastlines are suitable habitats for kelp forests^18,19^, but the actual extent of kelp forests is not fully mapped in most countries^31^ and is likely to exhibit seasonal and interannual variability in both extent and productivity^32^. Kelp-derived carbon stocks and fluxes (i.e., biomass, productivity, export, and sedimentary accumulation rates) leading to carbon sequestration are also uncertain because of natural variation and incomplete knowledge of their distribution, production, and POC and DOC fates. Moreover, not all exported kelp POC will be stored for long enough to be considered relevant for climate change mitigation (i.e., >100 years) and not all kelp POC that is stored will fall into existing carbon accounting and verification standards (i.e., within verifiable and governable reservoirs inside a country’s exclusive economic zone; EEZ)^14,15^. Given these uncertainties, new approaches are needed to estimate the current carbon sequestration capacity of kelp forests at national scales.

To facilitate accurate carbon accounting, we present a blueprint for producing national assessments of the blue carbon capacity of kelp forests (Fig. 1). Combining kelp data collation with Bayesian hierarchical modeling, this transparent and reproducible analytical framework estimates the carbon sequestration capacity of kelp forest ecosystems while explicitly acknowledging the inherent data limitations and uncertainties that most countries face in this regard. We apply this blueprint to Canada—a country accounting for 16.2% of the worlds coastline^33^–with expansive kelp forest ecosystems in the Atlantic, Pacific, and Arctic oceans.

**Figure 1.**
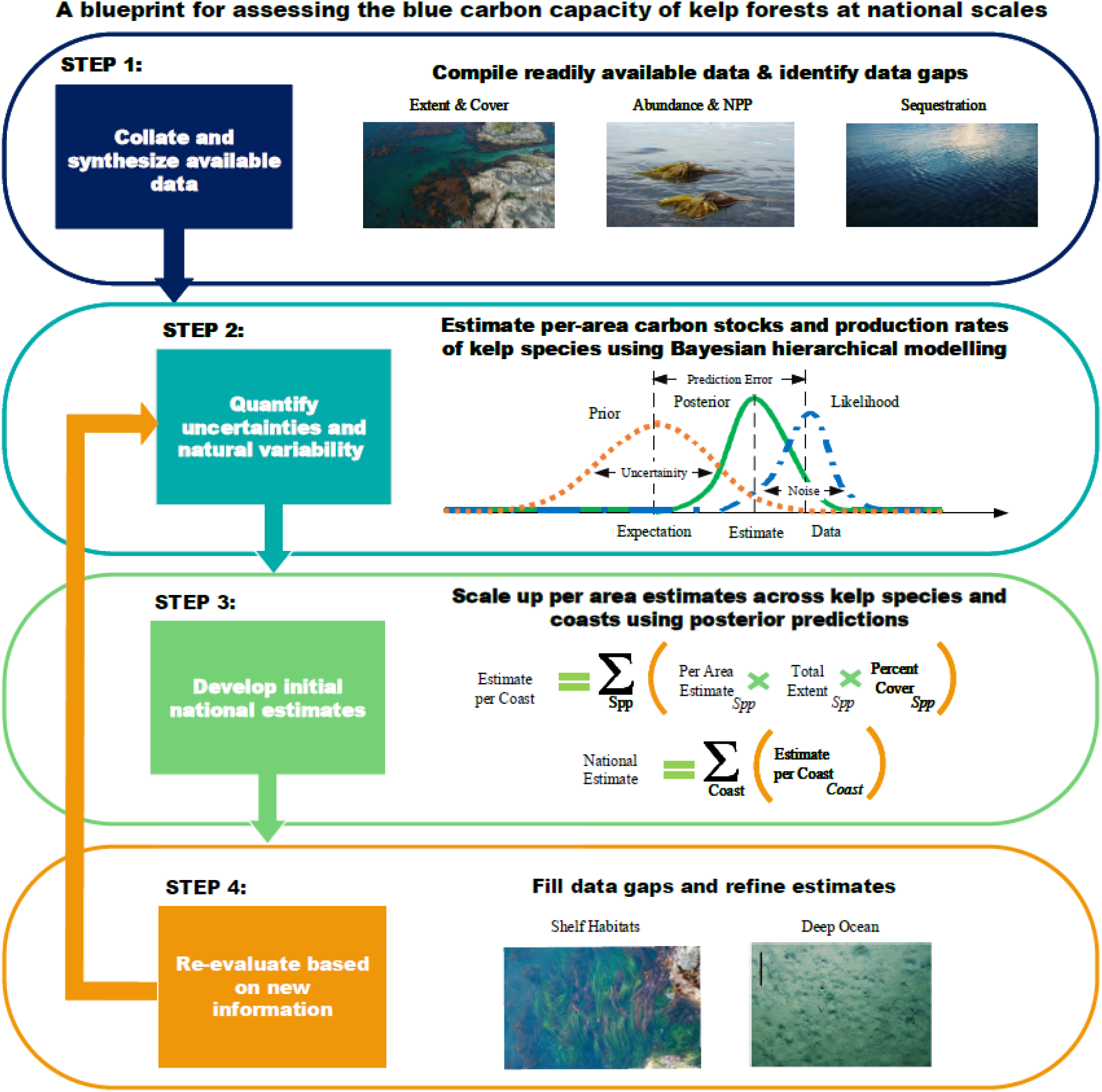
A proposed blueprint for national assessments of the blue carbon capacity of kelp forests. Our proposed blueprint involves steps to 1) compile and synthesize available kelp data, 2) quantify uncertainties and natural variability in potential rates of carbon production and storage by kelp species, 3) develop initial estimates of the carbon production, storage, and export capacity of kelp forests at national scales, and 4) refine assessments based on new information and data.

Two major surface canopy species, giant (*Macrocystis pyrifera*) and bull (*Nereocystis luetkeana*) kelp, form extensive floating forests along the Canadian Pacific, while subsurface species from the genera *Laminaria, Saccharina, Alaria, Agarum,* and others form dense submerged beds on their own, or as an understory below surface kelps, along substantial stretches of the Canadian coastline^40^. Our study enables the inclusion of kelp forest ecosystems into national NCS inventories in Canada and other countries with these important coastal ecosystems.

## Results

### Kelp forest blue carbon blueprint

Our blueprint for national assessments of the blue carbon capacity of kelp forests involves: 1) compiling and synthesizing available kelp data and identifying data gaps, 2) evaluating the potential for natural variation in the carbon stocks and carbon production rates for kelp species, 3) developing initial estimates of the standing carbon stock, production, and export capacity of kelp forests to deep ocean sinks, and 4) refining assessments based on new information and data (Fig. 1). For reproducibility, we provide a blueprint workflow and methodology for conducting an extensive collation of available datasets on the areal extent, canopy biomass, and NPP of kelp forests (Appendix A). We also provide R scripts that enable users to estimate the posterior mean carbon stocks and production rates of different kelp species based on limited available data and prior information using Bayesian hierarchical models (‘Brms’ package), as well as templates for scaling up per-area estimates to a national scale (Appendix B). Below we illustrate the blueprint’s utility through an application to Canada.

### First blueprint application: Canadian kelp forests

#### Data collation

We first compiled a database of kelp records from 36 published studies and monitoring programs (Appendix C: Table C1; Fig. C1) describing the areal extent, abundance (i.e., biomass and density), and NPP of subtidal kelp forest species across Canada’s Pacific, Atlantic and Arctic coasts (Fig. 1, Step 1). Our search targeted available data for surface kelp species found on the Pacific coast and subsurface kelps found across Canada’s three coasts, revealing that eleven of the 18 subtidal kelp species in Canada had sufficient data records to be included in further analyses. These include the two surface kelp species and seven of the 15 subsurface kelps located on the Pacific coast, five of the seven subsurface species found on the Arctic coast, and three of the five species found on the Atlantic coast (Table C2).

#### Kelp forest extents

##### Subsurface kelps

Next, since synoptic maps were unavailable, we produced high, mid, and low estimates of the potential extent of subsurface kelp forests in Canada using available depth, substrate, and kelp percent cover data (Table C3). To determine a hypothetical maximum limit for where subsurface kelp forests could occur in Canada, we calculated the area of rocky reefs (i.e., bedrock and boulders habitats) from mean-low-low-water out to 20 m water depth. With only this depth and substrate constraint, we found that subsurface kelp forests could cover up to 6.3 million hectares (Mha) (Table 1). Most of the kelp forest distribution (approximately 71%) was estimated to occur in the Arctic (5.5 Mha), while Atlantic and Pacific kelp forests covered 1.3 and 0.5 Mha, respectively (Appendix C: Table C3). Given that kelp do not always completely cover benthos, we then produced more constrained estimates for subsurface kelps, using available field surveys of kelp percent cover (Table C1), acknowledging that kelp percent cover can also vary annually and seasonally. We determined an upper biologically constrained extent of subsurface kelps by multiplying the maximum potential extent (described above) and the upper quartile of observed kelp percent cover at peak canopy biomass (May – August) across sites and years on each coast. We also determined a lower biologically constrained extent of subsurface kelp forests by multiplying the maximum potential extent by the lower quartile percent cover values, across years and sites, on each coast. Although the exact extent of subsurface kelp forests is still unknown, we estimated that the true extent falls between 0.8 and 3.9 Mha (Table C3).

**Table 1.** Comparison of the estimated extents, carbon stocks, carbon production rates, and carbon sequestration capacity of kelp forests, seagrass beds and salt marshes in Canada. Parenthetical values represent the lower and upper estimate values reported by this study and in the literature. *carbon sequestration for kelp forests is calculated in terms of the potential export of kelp detrital carbon to deep ocean sinks; carbon sequestration for seagrasses and salt marshes is calculated in terms of the amount of carbon accumulation in sediments. ND signifies no data for a particular field; NA signifies there the field is not applicable for a given ecosystem.

| Ecosystem | Areal extent (Mha) | C stock per-area (Mg C ha<sup>-1</sup>) | C production per-area (Mg C ha<sup>-1</sup> yr<sup>-1</sup>) | C sequestration per-area * (Mg C ha<sup>-1</sup> yr<sup>-1</sup>) | Standing C stock capacity (Tg C) | Standing C production capacity (Tg C yr<sup>-1</sup>) | Standing C sequestration capacity* (Tg C yr<sup>-1</sup>) |
| --- | --- | --- | --- | --- | --- | --- | --- |
| **Kelp forests** |  |  |  |  |  |  |  |
| *Canopy:* | 1.8<sup>+</sup><br>(0.8 – 6.3) | 0.8<sup>+</sup><br>(0.4 – 1.2) | 3.5<sup>+</sup><br>(1.3 – 6.7) | 0.6<sup>+</sup><br>(0.3 – 1.5) | 1.4<sup>+</sup><br>(0.6 – 4.6) | 3.1<sup>+</sup><br>(1.1– 11.6) | 0.2<sup>+</sup><br>(0.1 – 1.2) |
| **Seagrass meadows** |  |  |  |  |  |  |  |
| *Canopy:* | 0.8<br>(0.2 – 1.4)<sup>a</sup> | 0.1<br>(0.06 – 0.2)<sup>b</sup> | ND | NA | 0.08<br>(0.01 – 0.3) | ND | NA |
| *Soils:* | 0.8<br>(0.2 – 1.4)<sup>a</sup> | 88.2<br>(50.2 – 380.1)<sup>c</sup> | NA | 0.2<br>(0.04 – 0.9)<sup>a,c</sup> | 70.6<br>(10.0 – 532.1) | NA | 0.2<br>(0.01 – 1.3) |
| **Salt marshes** |  |  |  |  |  |  |  |
| *Canopy:* | 0.4<sup>d</sup> | ND | ND | NA | ND | ND | NA |
| *Soils:* | 0.4<sup>d</sup> | 80.4<br>(35.0 – 173)<sup>e</sup> | NA | 2.0<br>(0.6 -9.3)<sup>e</sup> | 32.2 | NA | 0.8 |

Data sources: <sup>+</sup>This study; <sup>a</sup>Drever et al. 2021, <sup>b</sup>Prentice et al. 2018, <sup>c</sup>Prentice et al. 2020 , <sup>d</sup>Rabinowitz & Andrews, <sup>e</sup>Kelly et al. 2023.

##### Surface kelps

As a particular case found on the Pacific Coast, we also produced high, mid, and low estimates specifically for surface kelp forests using available remote sensing and aerial surveys (Fig. C2). For the high estimate, we calculated the area of rocky reefs from mean-low-water out to 10 m water depth (Fig. C3). As an upper bound estimate, we used historical shoreline maps derived from oblique aerial survey imagery conducted by the British Columbia Shore Zone Survey from 2004-2007 to identify shallow rocky reefs that were previously covered by surface kelp forests. Finally, we used recent global surface canopy maps derived from Sentinel-2 satellite imagery from 2015 to 2019 as a low bound estimate. According to this analysis, surface kelp forests on the Pacific coast of Canada could cover up to 0.3 Mha, but more conservatively cover between 0.005 and 0.11 Mha (Table C3).

#### Per-area carbon stocks and production rates of kelp species

Bayesian hierarchical models revealed significant differences in per area carbon stocks and productivity within and among kelp species in Canada (Fig. 1, Step 2). On average, surface kelps tended to have higher values than subsurface species (Fig. 2). Giant and bull kelp, stored more carbon per area in their canopy biomass than six of the seven subsurface species (1.30 Mg C ha^-1^ and 0.95 Mg C ha^-1^, respectively), with over 80% conditional support for differences amongst the posterior mean predictions (Fig. 2a, Appendix C: Table C4). Giant and bull kelp also had the highest annual carbon production rates per area (7.26 Mg C ha^-1^ yr^-1^ and 6.35 Mg C ha^-1^ yr^-1^, respectively), producing more than twice the amount of carbon per year of other kelp species (Fig 2b; Table C4). While certain subsurface kelps (e.g., *Saccharina latissima*) had comparable estimated carbon stocks and production rates to surface kelps, most had much lower estimated carbon stocks per-area—ranging from 0.01 Mg C ha^-1^ (*Pleurophycus gardneri*) to 0.66 Mg C ha^-1^ (*Pterygophora californica)—*and carbon production rates per-area —ranging from 0.08 Mg C ha^-1^ yr^-1^ (*Agarum clathratum / Neoagarum fimbriatum*) to 3.18 Mg C ha^-1^ yr^-1^ (*Laminaria digitata / Hedophyllum nigripes*) (Fig. 2).

**Figure 2.**
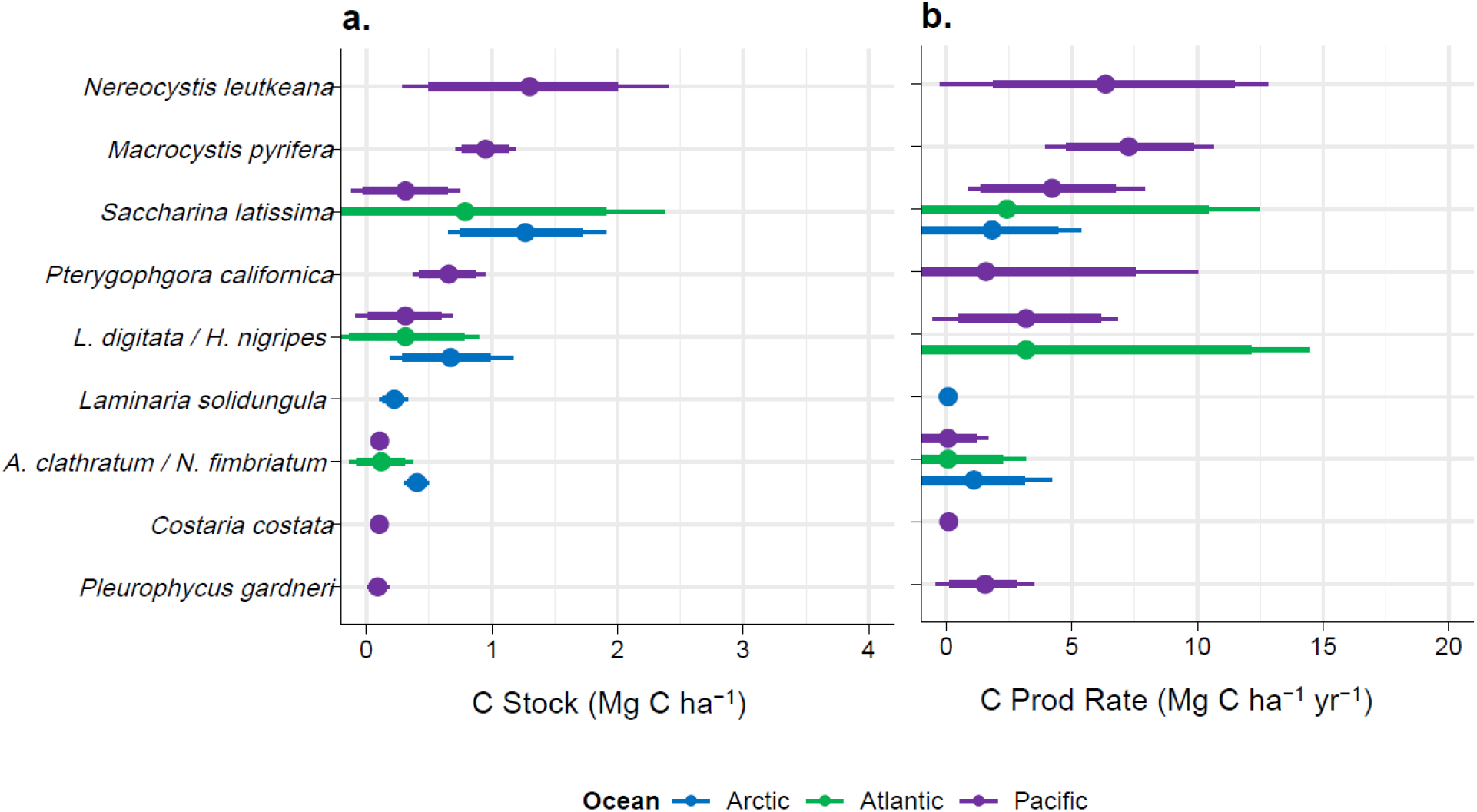
Per-area estimates of the a) carbon stock (Mg C ha^-1^) and b) carbon production (Mg C Ha^-1^ yr^-1^) capacity of kelp species across Canada’s three coastlines (Pacific = purple; Arctic = blue; and Atlantic = green) according to Bayesian hierarchical models. Posterior mean estimates (and 90% credible intervals) are shown for each species, representing the average posterior predictive distribution conditional on the observed data and prior information. The inner and outer bars show the credible intervals representing the range of values within which the true mean estimates are likely to occur with 80% and 90% probability based on the final models. Kelp species include: *Macrocystis pyrifera*, *Nereocystis leutkeana*,*Costaria costata, Agarum clathratum / Neoagarum fimbriatum*, *Laminaria digitata / Hedophyllum nigripes, Laminaria solidungula*, *Pterygophera californica*, *Pleurophycus gardneri,* and *Saccharina latissima*.

#### Per-area carbon stocks, production, and export rates of kelp forests by coast

Across Canada’s three coasts, we found considerable variation in the estimated per-area carbon stock and production rates of kelp forests due to differences in species composition and peak biomass (Fig. 3). Overall, Pacific kelp forests had the largest estimated carbon stocks per-area (1.2 Mg C ha^-1^) , along with the largest number of kelp species (N=17), and the highest estimated annual carbon production rates (6.7 Mg C ha^-1^ yr^-1^) (Fig. 3a). In comparison, Atlantic and Arctic kelp forests had lower kelp diversity (N=7 and 5, respectively) and a lower estimated carbon stock potential (0.4 and 0.8 Mg C ha^-1^, respectively), as well as much lower annual carbon production rates (2.7 Mg C ha^-1^ yr^-1^ and 1.3 Mg C ha^-1^ yr^-1^, respectively) (Fig. 3b).

**Figure 3.**
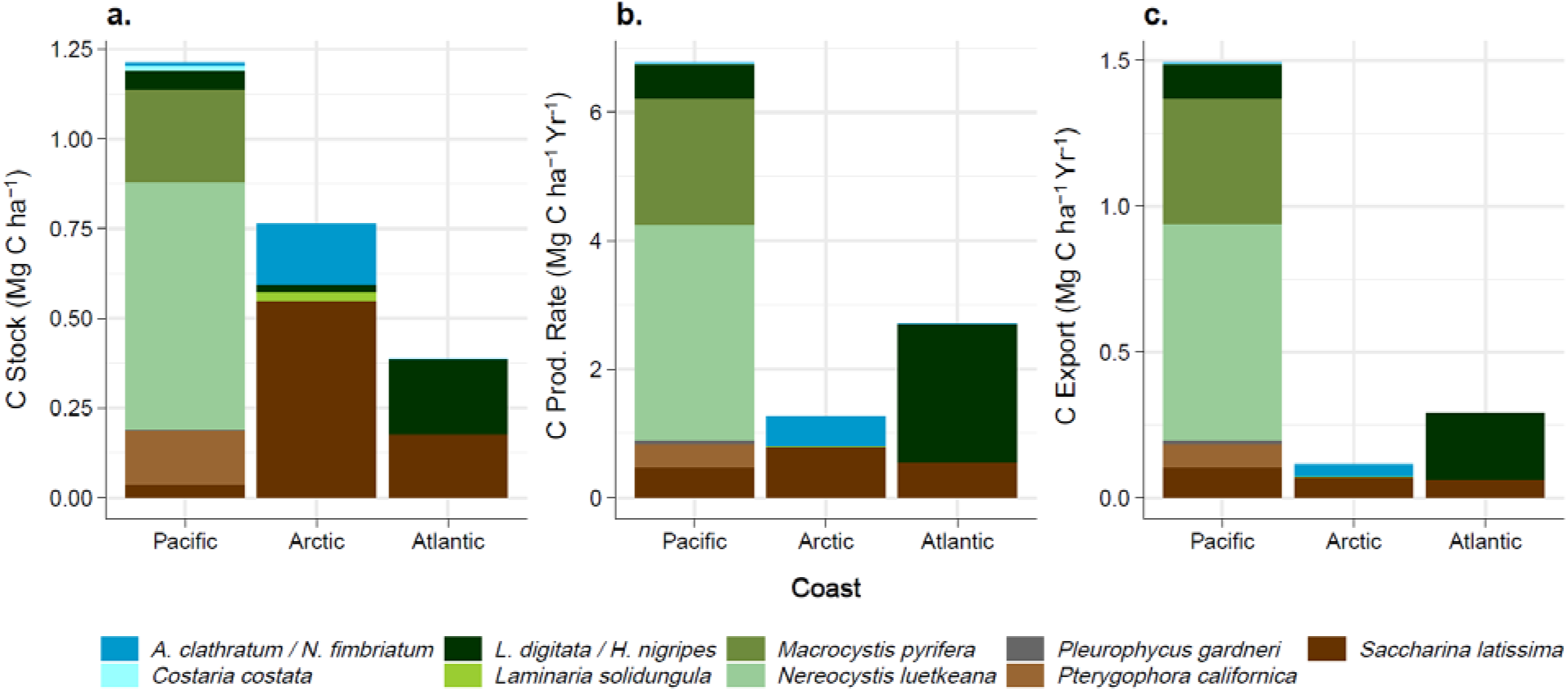
Per-area posterior mean estimates of the a) carbon stocks (Mg C ha^-1^), b) carbon production (Mg C ha^-1^ yr^-1^), and c) carbon export (Mg C ha^-1^ yr^-1^) capacity of subtidal kelp communities on Canada’s three coasts. Stacked bar plots show the summed posterior means across kelp species and coasts according to Bayesian hierarchical models, weighted by the relative abundance of kelp species on each coast.

As an approximation of the upper limit for carbon sequestration occurring in the deep ocean from Canada’s kelp forests, we also estimated the per-area rate of detrital export from kelp forests to beyond the continental shelf break (i.e., the 200-m isobath) according to global ocean transport estimates. Approximately 22.0% (SD = 12.0%) of kelp detritus is likely to reach the continent shelf break before decomposing in the Pacific coast compared to 10.8% (SD= 6.7%) in the Atlantic and 8.8% (SD= 2.8%) in the Arctic (Table B7). This implies that approximately 1.5 Mg C ha^-1^ yr^-1^ (0.7 – 2.3 Mg C ha^-1^ Yr^-1^) could be exported from Pacific kelp forests compared to 0.3 Mg C ha^-1^ yr^-1^ (0.2 – 0.4 Mg C ha^-1^ yr^-1^) from Atlantic kelp forests and 0.1 Mg C ha^-1^ yr^-1^ (0.01 – 0.2 Mg C ha^-1^ yr^-1^) from Arctic kelp forests (Fig. 3c).

#### First national estimates for Canada’s kelp forests

Finally, to produce national estimates of the carbon sequestration capacity associated with Canada’s kelp forests, we combined the kelp forest extent estimates with the per-area carbon stock, carbon production, and carbon export estimates on each coast (Fig. 1, Step 3). For a conservative scenario, assuming kelp forests are at their median areal extent, we estimate that Canadian kelp forests have a standing carbon stock capacity of 1.4 Tg C (0.6 - 2.8 Tg C) and an annual carbon production capacity of 3.1 Tg C yr^-1^ (1.1 – 6.3 Tg C yr^-1^), approximately 0.2 Tg C Yr^-1^ (0.04 – 0.4 Tg C yr^-1^) of which could be transported to and sequestered in the deep ocean (Fig. 4). However, in the most optimistic scenario, where kelp forests are at their maximum potential extent, these figures increase to a national standing stock capacity of 4.4 Tg C, an annual carbon production capacity of 11.6 Tg C yr^-1^, and an annual carbon export capacity of 1.0 Tg C yr^-1^ to the deep ocean. Arctic kelp forests had the greatest overall carbon stock (1.3 Tg C; 0.6 – 3.5 Tg C) and production capacity (2.1 Tg C yr^-1^; 1.0 – 5.8 Tg C yr^-1^) (Fig. 4a - b) because of their disproportionately larger areal extents. However, kelp forests in the Pacific had the highest estimated capacity for carbon sequestration via export to the deep ocean (0.15 Tg C yr^-1^, 0.01 – 0.5 Tg C yr^-1^) because of the higher per-area carbon production rates and potential for detrital transport beyond the shelf break (Fig. 4c).

**Figure 4.**
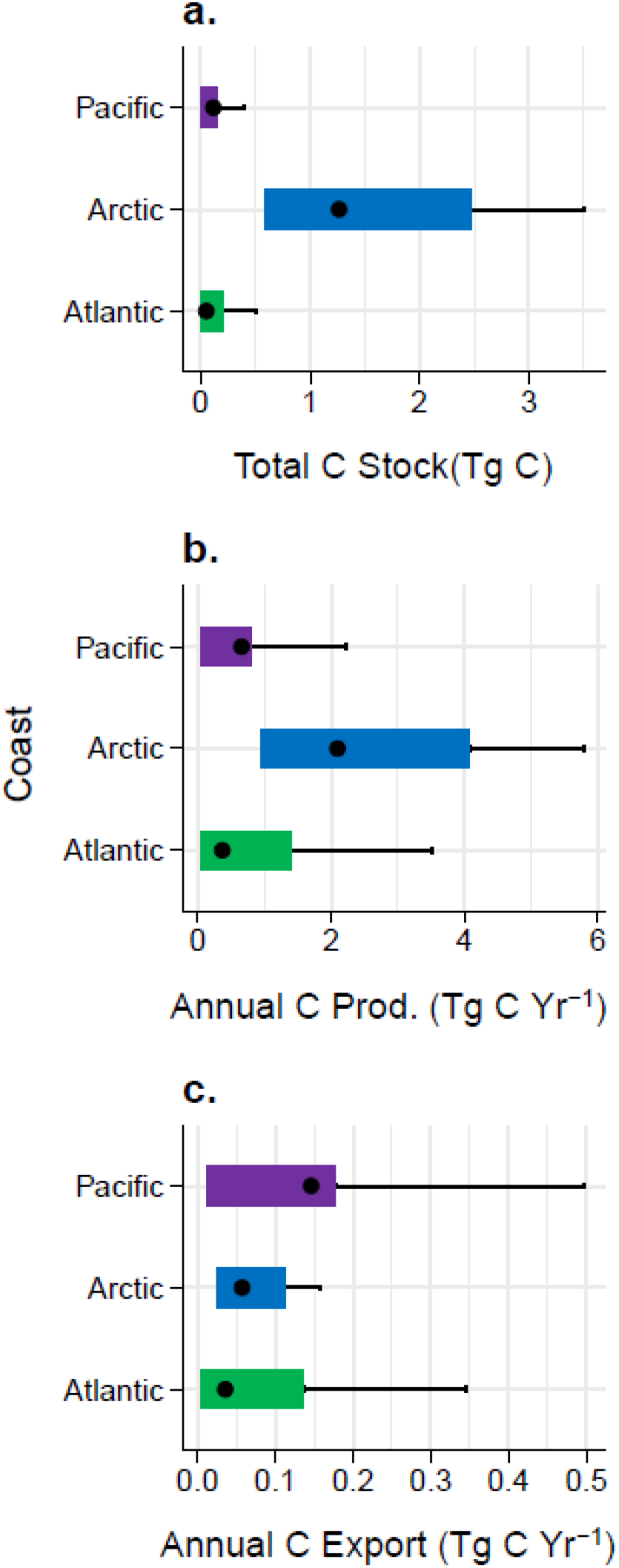
National carbon capacity of Canadian kelp forests depicted in terms of the total estimated a) standing carbon stocks (Tg C), b) carbon production (Tg C yr^-1^), and c) carbon export (Tg C yr^1^) capacity of kelp forests. The bars depict the upper bound (75^th^ percentile) and lower bound (25^th^ percentile) estimates per coast. The circle represents the median estimates for each coast. Error bars show the maximum potential capacity per coast.

## Discussion

National assessments of BCEs, such as seagrasses, salt marshes, and mangroves, are becoming more prevalent^2,34^, paving the way for their incorporation into NCS inventories. However, comparable evaluations for kelp forests are currently unavailable for nearly 90% of the 150 countries with kelp forests^14^, due to existing data gaps and the difficulty of accurately estimating the kelp-derived carbon stocks and fluxes leading to sequestration in various ocean sinks (i.e., DOC pools, shelf sediments, and the deep ocean). Our reproducible blueprint, applied to Canadian kelp forests, has important implications for other countries looking to account for kelp forests as NCS.

### Potential for Canada’s kelp forests as an NCS

Our assessment found the carbon production capacity of Canadian kelp forests to be substantial (3.1 Tg C yr^-1^; 1.1 - 11.5 Tg C yr^-1^). This figure is low compared to recent global estimates of kelp carbon production (∼1.5% of global estimated NPP) but these are not directly comparable as we used a more conservative depth cut-off when calculating the extent of kelp forests (20m compared to 30m)^18^. Additionally, we found that Canadian kelp forests provide a clear pathway for sequestering and storing carbon in the deep ocean (0.2 Tg C yr^-1^, 0.04 to 1.0 Tg C yr^-1^). Realised carbon sequestration of kelp-derived carbon could be even greater when accounting for kelp carbon entering refractory DOC pools in the deep ocean. Compared to terrestrial ecosystems, kelp forests are likely to play a more modest role in the overall NCS components of Canada’s climate change mitigation strategy^3^. As examples, conservation pathways for grasslands, peatlands, and forests have been estimated to sequester 3.5 Tg C Yr^-1^ , 2.8 Tg C yr^-1^, and 2.2 Tg C yr^-1^, respectively^3^. Nevertheless, we found that kelp forests could have comparable carbon sequestration benefits to freshwater mineral wetlands (0.8 Tg C yr^-1^)^3^ and other BCES, such as eelgrass meadows (0.2 Tg C yr^-1^), and tidal marshes (0.8 Tg C yr^-1^) (Table 1), thus warranting further consideration in Canada’s NCS inventories.

We found contrasting patterns of carbon production, storage, and export across Canada’s coastlines, reflecting different kelp species assemblages and environmental conditions across these vast areas. Notably, the per-area carbon production capacity of Pacific kelp forests exceeded the global averages for subtidal kelps, intertidal seaweeds, subtidal red seaweeds, and floating seaweeds (e.g., *Sargassum* spp)^18^. While the Arctic had the highest total kelp standing carbon stock and production capacity, due to their extensive coastline and wide continental shelf, Pacific and Atlantic coasts had a higher capacity for kelp carbon sequestration in the deep ocean overall, because of their higher per-area rates of kelp carbon production and hydrological export. However, kelp forests on all three coastlines showed some potential to sequester carbon in the deep ocean, highlighting their potential role in Canada’s NCS inventories.

Our assessment revealed considerable data gaps across all elements of our analysis, underscoring the need for kelp monitoring programs at a national scale. Given the lack of comprehensive habitat maps, we needed to make assumptions about the current extent of kelp forests, including assumptions about the maximum depth limit of kelp forests, the prevalence of rocky reefs, and the kelp occupancy and abundance across Canada’s coastline. We also could not account for ecological driver (e.g., urchins) that likely limit kelp extents in certain areas^35^. When estimating per-area carbon stocks and production rates, we faced significant data limitations for many kelp species, especially the subsurface kelps, leading to large credible confidence intervals for many species. Additionally, we needed to make assumptions about the relative abundance of species in kelp forests when extrapolating standing carbon stock, production, and export estimates to the coast-wide scale. Lastly, given the complete lack of data on the accumulation of kelp derived carbon in shelf and deep ocean sinks, we relied on hydrological export estimates from ocean transport models to approximate carbon sequestration rates beyond the continental shelf breaks (as only one potential pathway of kelp carbon sequestration). A sensitivity analysis revealed that the maximum depth limit and the hydrological export rates are likely to have the strongest influence on national estimates (Fig. C4 – C6), suggesting these datasets should be the highest priority for future research. However, greater investment in collecting of kelp species abundance, composition, and net primary productivity data is also needed to estimate kelp carbon sequestration more accurately nationally. Notably, many of these data can be and are already being collected by coastal communities and First Nations in Canada (e.g., through the Marine Plan Partnership program)^36^, creating future opportunities for collaborative research efforts.

### General considerations of kelp NCS

While highlighting kelp forests in Canada as a potential asset for storing and sequestering atmospheric carbon, our study has broader implications for developing NCS in other data-limited countries with kelp forests. First, our findings underscore the importance of elucidating and considering the full pathways of carbon production, export, and storage. For example, despite the seemingly higher total carbon stocks and production capacity of Canada’s Arctic kelp forests, a comprehensive understanding of the area-based production and export potential of distinct coastlines was needed to reveal the greater contributions of Pacific and Atlantic kelp forests to carbon export fluxes to the deep ocean. However, it is possible that Arctic kelp forests could play a more important role in shelf carbon sequestration, particularly given the Arctic’s expansive shallow continental shelf and the greater potential for preservation due to cold temperatures^37^. Additionally, our findings emphasize the potential importance of spatial and temporal variation in kelp carbon cycling. In Canada, potential export rates varied by an order of magnitude difference(0.9 to 33.6%)depending on the ecoregion^38^. Additionally, kelp species showed considerable variation in their estimated carbon stocks and production rates, which may increase further as more spatially and temporally resolved data becomes available. Overlooking these variables could lead to biased estimates, potentially undermining the effectiveness of NCS.

Our findings imply that continued environmental changes could have varying consequences for kelp carbon sequestration. Kelp degradation and deforestation have occurred globally due to various anthropogenic stressors and disturbances, including overfishing, eutrophication, climate change and species invasions^39–43^. For instance, along Canada’s Pacific and Atlantic coast, kelp declines have been documented after unchecked urchin grazing and intensifying marine heatwaves^35,44^, while many kelp forests in Atlantic Canada have transitioned to beds dominated by algal turfs due to the combined effects of warming temperatures and interactions with invasive species^40,45,46^. These changes are likely to have severe consequences for associated biodiversity and other ecosystem functions (e.g., fisheries production), and they may also disproportionately reduce the capacity of kelp forests to produce and export carbon.

Variation in the response of different kelp species to climate change may also have important implications for understanding the impacts of kelp species redistribution on carbon sequestration^47^. As kelp distributions are altered by warming temperatures, there could be considerable changes in kelp community composition^48,49^; additionally, more frequent marine heatwaves may lead to local extirpations of kelp species, which could impact carbon production and storage patterns^32,50^. It is possible these changes could lead to enhanced kelp carbon sequestration at the cold edge of species’ ranges. For instance, models from the Arctic show the possibility of range expansions for *S. latissima, A. clathratum,* and *A. esculenta* with the loss of sea ice and warming ocean temperatures^51^, which could further increase overall carbon production in this region. However, warming ocean temperatures could also lead to faster decomposition rates^52^, and it is unknown whether these gains in suitable area and productivity would offset losses occurring at the warmer range edges^53^ or locally warm hotspots^32,40,54^. Ultimately, expanded monitoring datasets and better forecasting models are needed to understand the full scope of climate impacts on kelp carbon sequestration.

### Applying the blueprint

Our blueprint can be applied to other countries, providing a roadmap for evaluating carbon stock, production, and export capacity in other kelp-dominated systems. Developed in a country with an with extensive coastlines, diverse kelp communities, and complex oceanographic settings, our approach is useful for evaluating kelp forests wherever there is data on the areal extent, abundance, and net primary productivity of kelp species (see Fig. 1). For coastal countries where comprehensive maps of kelp forest extent are not yet available, coarse approximations could be obtained using global data on coastal bathymetry and existing global species distribution models^18,50^. Additionally, publicly available data on the NPP of kelps can be extrapolated from other systems and global models, and used as prior information and data in the absence of regional datasets^55^. In the absence of regional models, empirical measurements of rates of kelp carbon export can be supplemented with ocean transport models, which can help approximate coastal to open ocean transport to various long-term sinks (e.g. ^23,56^). However, current export models will require a thorough interrogation with *in-situ* experiments and observation studies.

Through integrating Bayesian hierarchical models with extensive data collation and synthesis, our approach addresses prevailing challenges associated with estimating species-specific carbon stocks and productivity rates, including data scarcity and the unknown potential for variability. One key advantage lies in the ability of Bayesian models to leverage prior information about the known range and variability of kelp productivity from related species and systems when making posterior predictions. Bayesian hierarchical models can also allow for incorporating different forms of measurement error (e.g., standard deviations in field measurements across years) for a more transparent accounting of the residual uncertainty. Furthermore, our approach produces national scale estimates in terms of a conservative range from a lower bound to a maximum potential as determined by prior information and data. By providing this range, our approach acknowledges the inherent complexities and variability of kelp ecosystem dynamics by providing a range estimate, offering decision-makers a more comprehensive and yet cautious perspective to guide policy and management strategies.

### Priority Research Directions

Application of our blueprint to Canada’s kelp forests underscores general uncertainties and priority research directions for countries attempting to incorporate kelp forests in their NCS inventories. First, our assessment estimates the standing carbon production capacity of kelp forests, from which the amount of kelp POC predicted to reach the continental shelf break each year can be used as a proxy for carbon fluxes to the deep ocean. However, the magnitude of kelp primary production that will be exported, sequestered, and stored in the deep ocean will depend on relative rates of degradation, vertical exchange, sediment accumulation, sediment remineralization, and other factors^20,29,37,57^. Additionally, the export of kelp POC to the deep ocean is just one potential pathway of carbon sequestration (e.g., POC fluxes to other BCEs and DOC fluxes to the deep ocean). Acquiring more comprehensive datasets on carbon accumulation, export, and retention rates across various reservoirs is crucial for refining estimates of the carbon sequestration benefits from these important ecosystems.

Second, significant questions remain about whether carbon sequestration by kelp forests could lead to meaningful climate change mitigation benefits either through avoided emissions or restoration pathways^14,25^. Demonstrating the impact of kelp forests on air-sea CO_2_ fluxes is challenging and whether air-sea fluxes can offset local respiration rates in kelp forests remains uncertain^25,58^. Additionally, for kelp forests to meaningfully contribute to climate change mitigation, interventions must modify GHG emissions and/or removals beyond what would happen naturally (i.e., additionality). Proposed interventions, including protection of threatened kelp forests, restoration of lost kelp forests, and artificial expansions of kelp forests beyond their historical extents via seaweed mariculture^30^ would need to create durable GHG gains (i.e., >100 years) and fall within a country’s jurisdictional boundary to be eligible for policy and management action^15^. Comparing estimates of maximum potential carbon sequestration with projections based on actual kelp distribution data could provide valuable insights into opportunities for further enhancement. However, further research is needed to understand the full scope of options for kelp based NCS.

Finally, as we move towards a future characterized by ocean warming and intensifying marine heatwaves, there is a pressing need to understand how these changes could impact the carbon sequestration capacity of kelp forests. Improved forecasting models and expanded monitoring efforts are essential to anticipate changes in kelp carbon sequestration and to develop climate-smart management. Integrating kelp forests into national and global climate change mitigation strategies also requires robust and standardized methodologies for quantifying and verifying carbon stocks and fluxes to ocean carbon sinks under future scenarios^59^.

## Conclusions

As nations strive to meet their net zero targets in carbon emission, incorporating kelp forests into NCS inventories represents an important next step in harnessing the full potential of BCEs. To that end, countries must be able to reliably estimate and predict changes in kelp carbon sequestration resulting from proposed management, conservation, and restoration actions. Our analytical framework offers a blueprint for developing initial assessments across a range of systems, representing a significant step forward for blue carbon accounting for kelp forests. Applying this framework to Canada, we show that estimated annual fluxes of kelp-derived carbon to the deep ocean are at least comparable to the carbon sequestration capacity of other BCEs in Canada, suggesting that kelp forests merit further consideration within Canada’s GHG inventories. Additionally, our study highlights many of the important considerations, data needs, and uncertainties surrounding the accounting of kelp carbon sequestration at national scales. Our study can serve as an important resource for policymakers, researchers, and stakeholders aiming to integrate kelp forests into their climate action strategies.

## Methods

### Study area

Our study area spans the Pacific, Arctic, and Atlantic coasts of Canada, from mean sea level out to the 20-meter depth contour. In the Pacific, this includes 25,000 km of coastline from 48 to 55° N; in the Arctic, 162,000 km from 51 to 83° N; and in the Atlantic, 42,000 km from 43 to 60° N. These coasts support a diversity of kelp forest-forming species. Indeed, the Northeast Pacific Ocean is considered the evolutionary center of origin for kelps^60^ and includes >30 kelp species that vary in morphology and ecological niche^61^. The Arctic and North Atlantic oceans were subsequently colonized and recolonized following glaciation events and are now home to >10 kelp species^62,63^.

### Data scoping

We collated information and datasets from a variety of published and unpublished sources on the areal extent, biomass, plant density, canopy cover, and NPP of the most common kelp forest species (Table C1). We limited our search to all surface and subsurface kelp species found in the subtidal zone of at least one Canadian coast, according to global species-occurrence databases^19,64^. To collate sources from the published literature, we used an existing database of macroalgal NPP measurements compiled from a combination of reports, peer-reviewed studies, and PhD and masters theses published between 1967 and 2021^55,65^. Following similar search criteria, we expanded this NPP database to include additional papers published between 2021 – 2023 for Canadian kelp species. We then used this updated NPP database to compile published measurements of kelp biomass from the text, figures, and supplementary datasets of the original source material. In addition, we compiled datasets from unpublished sources using a snowball search method, where we reached out to the authors of previously published kelp papers in Canada and asked for recommendations on potential data sources for kelp extent, biomass, canopy cover, and net primary productivity.

We focused subsequent analyses on the kelp species that had at least one biomass and NPP record for a species on a given coast (Table C2). These included the two surface kelp species (i.e., *Macrocystis pyrifera* and *Nereocystis luetkeana*) and seven of the 15 subsurface kelps found on the Pacific coast (*i.e., Agarum clathratum, Costaria costata*, *Hedophyllum nigripes*, *Neoagarum fimbriatum, Pterygophora californica, Pleurophycus gardneri, Saccharina latissima*); five of the seven species found on the Arctic coast (i.e., *A. clathratum, Laminaria digitata, L. solidungula, H. nigripes*, and *S. latissima*; all subsurface); and three of the five species found on the Atlantic coast (*i.e., A. clathratum, L. digitata,* and *S. latissima*; all subsurface). Since *H. nigripes* could not be differentiated from *L. digitata* in some of the Arctic and Atlantic data records, the two species were grouped together in subsequent analyses. Likewise, we grouped *A. clathratum* and *N. fimbriatum* records due to the difficulties with differentiating these two species in the field.

### Determining the potential extent of kelp forests

We produced high, mid, and low estimates of the potential areal extent of subsurface kelp forest in Canada using available depth, substrate, and kelp percent cover data. As a hypothetical maximum potential extent for subsurface kelps, we calculated the area of suitable rocky seafloor across Canada, i.e., the areal extent of rocky seafloor between mean low water and the 20 m depth contour in millions of hectares (Mha)^50^. This is a conservative depth cutoff since kelp forests occur deeper (50 m) in some areas^66^. Depth estimates were based on the General Bathymetric Chart of the Ocean data (https://www.gebco.net; GEBCO)—a gridded global terrain model for the ocean and land at 15-arc-second resolution. The extent of rocky seafloor was based on public spatial data repositories for the Pacific and Atlantic coasts^67,68^. Since there was limited information on the distribution of rocky seafloor for most of the Atlantic coast, we used the fraction of rocky seafloor found on the Scotian shelf (30.7 %) as a coarse proxy^67^. For the Arctic, we used the fraction of rocky seafloor used by previous global studies (20%)^18^. Finally, we masked extents in the Arctic by the areal coverage of perennial sea ice occurring at the northern edges, using spatial data layers from BioOracle^69^.

To constrain the upper and lower bound estimates for the potential extent of subsurface kelp forests, we combined the maximum potential extent maps with existing field surveys of the percent cover of kelp forests from the Pacific, Atlantic and Arctic coasts. We acquired quadrat surveys of the percent cover of subsurface kelp species from active monitoring programs^70,71^ and the peer-reviewed literature (Table C1). To calculate the upper bound extent of subsurface kelp forests, we multiplied the maximum potential extent by the 75^th^ percentile of observed percent cover for all kelp species (regardless of the species composition) on each coast.

Additionally, we calculated the lower bound as the maximum potential area multiplied by the 25^th^ percentile of observed kelp cover.

We also produced high, mid, and low bound estimates for the potential areal extent of surface kelp forests, as a particular case solely found on the Pacific coast of Canada. First, we calculated the maximum potential extent of surface kelps as the area of suitable rocky seafloor above 10m water depth—the depth above which 90% of the observations for bull kelp and giant kelps occur in British Columbia (Fig. C3). To produce mid and low bound estimates for surface kelps, we used available coarse maps derived from existing aerial- and satellite-remote sensing products. We determined the shoreline distribution of *M. pyrifera* and *N. luetkeana* according to their presence in georeferenced oblique aerial imagery collected and analyzed by the British Columbia Shore Zone program from 2004 to 2007 ^72,73^. From this dataset, we calculated the upper bound of potential surface kelp extent as the intersection between previous shoreline detections (i.e., within 500 m) and the maximum area of suitable rocky seafloor adjacent to the shoreline. Finally, we acquired global distribution maps of surface canopy kelps determined from classified 20m resolution Sentinel-2 satellite imagery from 2015 to 2019^74^, which we used as the lower bound extent of surface canopy forming species. We validated both datasets through expert comparison with Google Earth Imagery, removing obvious false positives found in higher estuaries and along the intertidal zone, in ArcGIS Pro Version. 3.0.

### Determining the per-area carbon stocks of kelp species

We quantified carbon standing stocks associated with kelp forests in Canada by compiling available data on the area-specific biomass and plant density of kelp species from published and unpublished sources (Table C1). Wet weight measurements (i.e., g WW m^-2^ ) for each species and coast were converted to dry weight (g DW m^-2^) using species- and coast-specific conversions from the peer-reviewed literature ^55^. Kelp densities (i.e., number m^-2^) for each species and coast were also converted to dry weight using available average individual weight measurements ^55^. We then used species- and coast-specific ratios to convert dry weight measurements to the area-specific organic carbon content (g C m^-2^) of each sample ^55^. Finally, we converted all measurements of organic carbon content to carbon standing stocks in units of megagrams per hectare (Mg ha^-1^) by species.

### Determining the per-area annual carbon production rates of kelp species

We used available published and unpublished measurements of net primary productivity for all kelp species found in Canadian waters (Table C1)^55,65^. We also used published net primary productivity records from locations with similar environmental conditions to Canadian waters (i.e., within the range of mean ocean temperatures observed on the Pacific, Arctic, and/or Atlantic coasts according to BioOracle data layers)^69^. All wet weights were converted to dry weight measurements and then all dry measurements were converted to area-specific rates of net primary productivity (i.e., g C m^-2^ yr^-1^), using species- and coast-specific conversions from the literature^55^. We then converted all measurements of NPP to annual carbon production in units of megagrams per hectare per year (Mg ha^-1^ Yr^-1^) by species.

### Bayesian hierarchical models

We used Bayesian hierarchical models to evaluate the potential for natural variation in the per-area carbon stocks (Mg C ha^-1^) and carbon production rates (Mg C ha^-1^ yr^-1^) of different kelp species in Canada. Bayesian hierarchical models are parameterized similarly to hierarchical linear regression models using the Stan computational framework (http://mc-stan.org/), which can be accessed via the ’brms’ package of the R programming language (version 2022.12).^75^. A key advantage of Bayesian hierarchical models is the ability to generate a posterior distribution, representing the central tendency (i.e., the posterior mean) and the probabilistic range of uncertainty surrounding a parameter estimate. A credible confidence interval (CCI) can be derived from this posterior distribution, indicating the range within which the true parameter value is likely to fall. Additionally, the ‘brms’ package allows for the explicit consideration measurements of standard deviation as an additional response term, allowing for adjusted CCIs that reflect greater uncertainty where there is greater variability in kelp biomass and NPP. Finally, a Bayesian approach allows for the use of informative priors based on observations from different systems that further constrain the parameter estimates and CCIs. Additional information about using the R package “brms” can be found in the literature and the documentation^76,77^. Scripts for model parameterization, selecting informative priors, and evaluating model outputs, can also found in our Github repository (https://github.com/jennmchenry1/A-blueprint-for-national-assessments-of-blue-carbon-capacity-of-kelp-forests-CA).

The observed carbon standing stocks and production rates of eleven kelp species were modeled as the response variables. To account for measurement uncertainty in the available observations, we included the standard deviation measurements representing the site-level variation as an additional response term. For each species, we built sets of competing models that tested the effects of different combinations of predictors on species carbon stocks and production rates (Table C5). We accounted for potential fixed effects of mean sea surface temperatures derived BioOracle^69^ and the oceanic context (i.e., Pacific, Arctic, and Atlantic), and controlled for the sampling year and site identity as random effects. The models ran for 5000 iterations with 2500 warm-ups using three chains. Convergence was visually assessed by examining the trace plots and further verifying all coefficients achieving an Rhat value of 1^78^. To determine which model best described each response variable, we used an approximation for leave-one-out (LOO) cross-validation (‘loo’ package)^79,80^. We evaluated the performance of the final models through a series of posterior predictive checks where draws from the posterior distribution of model parameters were compared to the observed data as a measure of model goodness of fit (Fig. C7 – C8).

Models were trained with weakly informative ‘priors’, setting the scale of the prior distribution to be larger than and consistent with the range of potential observed values in our collated response datasets (Table C6) and the range of global synthesized primary productivity measurements from macroalgal forests ^65^. To ensure that our choice of priors did not overly constrain the resulting posterior predictions or inflate the uncertainty intervals, we conducted a prior sensitivity analysis for the three most data rich species in our dataset (i.e., *M. pyrifera, N. leutkeana*, and *S. latissima*) and used the best matched set of priors for the remaining species.

We present the final models for eleven kelp species, which were selected by the approximate LOO cross-validation (Table C7). The final models were used to generate posterior mean estimates of the potential carbon stocks and production rates associated with kelp species, including the 90% credible confidence intervals around those estimates. Significant differences among the posterior mean estimates were assessed through comparison of percent overlap between credible confidence intervals.

### Estimating the national carbon stocks, production, and sequestration capacity of kelp forests

We estimated the standing carbon stocks (Tg C) of current kelp forests in Canada as the summed product of the kelp forest extent (E_coast_) and the carbon stock potential of kelp forests across Canada’s three coastlines (CStock_coast_) (Equation 1). As inputs to this calculation, we used the posterior mean estimates of the carbon stocks of individual kelp species (described above). To account for the fact that kelps often persist in multi-species assemblages and thus are not likely persisting at their maximum biomass potential, we estimated the per-area carbon stock of kelp forests per coast (CStock_coast_) as the summation of the posterior mean estimates for each kelp species (CStock_spp_), weighted by the relative abundance of that kelp species (A_spp_), on each coast. We used the maximum, upper, and low kelp forest extent estimates as inputs to determine the most likely maximum, upper bound, and lower bound carbon stock potential of each coast.

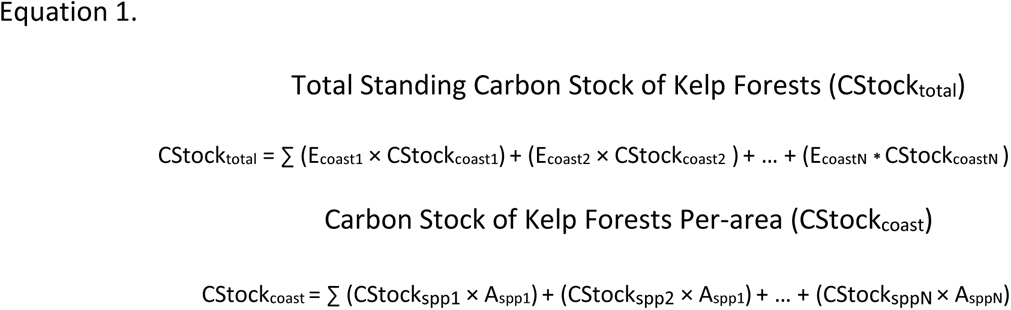

Additionally, we estimated the total annual carbon production capacity of kelp forests (Tg C yr^-1^) of current kelp forests in Canada as the summed product of the kelp forest extent (E_coast_) and the carbon production rate of kelp forests across Canada’s three coastlines (CProd_coast_) (Equation 2). To estimate the per-area carbon production rate of kelp forests per coast (CProd_coast_), we summed the posterior mean estimates for each kelp species (CProd_spp_), weighted the relative abundance of that kelp species (A_spp_), on each coast. We calculated the total carbon production capacity of kelp forests per coast in terms of the maximum, upper bound, and lower bound extent estimates.

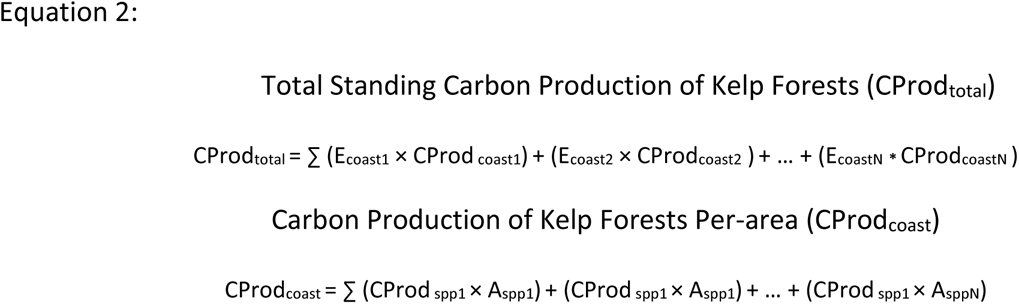

Finally, we estimated the total annual capacity (Tg C yr^-1^) for the export of kelp-derived carbon beyond the continental shelf break (i.e., the 200-m isobath), as an approximation for the total carbon sequestration in the deep ocean resulting from Canada’s kelp forests (Equation 3). To do so, we acquired estimates of the fraction of kelp carbon detritus (Exp_ecoregion_) that may be exported to the open ocean before decomposing according to a global model of shelf to open ocean exchange for all ecoregions falling within Canada’s EEZ^38^. We determined the total annual export capacity of kelp forests in Canada as the summed product of the estimated kelp forest extent (E_coast_) and the annual carbon export rate of kelp forests across Canada’s three coastlines (CFlux_coast_) (Equation 3). As an input, we estimated CFlux_coast_ as the summation of the fraction of modeled hydrological export per ecoregion (Exp_ecoregion_) multiplied by the per-area annual carbon production of kelp forests for a given coast (CProd_coast_; calculated above). We calculated the total carbon export capacity of kelp forests per coast in terms of the maximum, upper bound, and lower bound extent estimates.

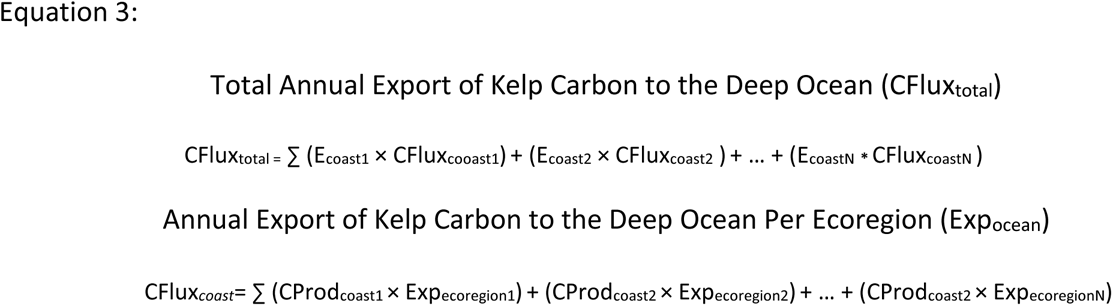

## Acknowledgements

We acknowledge funding and research support from Fisheries and Oceans Canada (*contract number: 6861*), the National Science and Research Council of Canada’s Alliance grant program (*grant number: ALLRP 571068-21*), the Mitacs Accelerate grant program (*grant number: IT28407*), The Tula Foundation and the Hakai Institute, ArcticNet (*ArcticKelp P101*) and the Australian Research Council (*DP220100650*).

## Data Availability Statement

All research outputs, including the collated and synthesized datasets and the resulting national estimates, will be made available at the time of publication through the data repository, Borealis. Additionally, the R code supporting this study will be uploaded to a Github repository (https://github.com/jennmchenry1/A-blueprint-for-national-assessments-of-blue-carbon-capacity-of-kelp-forests-CA).

## References

1. Griscom, B. W. et al. Natural climate solutions. Proc. Natl. Acad. Sci. 114, 11645–11650 (2017).

2. Macreadie, P. I. et al. Blue carbon as a natural climate solution. Nat. Rev. Earth Environ. 2, 826–839 (2021).

3. Drever, C. R. et al. Natural climate solutions for Canada. Sci. Adv. 7, eabd6034 (2021).

4. Fargione, J. E. et al. Natural climate solutions for the United States. Sci. Adv. 4, eaat1869 (2018).

5. Alongi, D. M. Global Significance of Mangrove Blue Carbon in Climate Change Mitigation. Sci 2, 67 (2020).

6. Chmura, G. L., Anisfeld, S. C., Cahoon, D. R. & Lynch, J. C. Global carbon sequestration in tidal, saline wetland soils. Glob. Biogeochem. Cycles 17, (2003).

7. Fourqurean, J. W. et al. Seagrass ecosystems as a globally significant carbon stock. Nat. Geosci. 5, 505–509 (2012).

8. Mcleod, E. et al. A blueprint for blue carbon: toward an improved understanding of the role of vegetated coastal habitats in sequestering CO _2_. Front. Ecol. Environ. 9, 552–560 (2011).

9. Bufarale, G. & Collins, L. B. Stratigraphic architecture and evolution of a barrier seagrass bank in the mid-late Holocene, Shark Bay, Australia. Mar. Geol. 359, 1–21 (2015).

10. Duarte, C. M., Middelburg, J. J. & Caraco, N. Major role of marine vegetation on the oceanic carbon cycle. Biogeosciences 2, 1–8 (2005).

11. Lo Iacono, C., et al. Very high-resolution seismo-acoustic imaging of seagrass meadows (Mediterranean Sea): Implications for carbon sink estimates. Geophys. Res. Lett. 35, (2008).

12. Waycott, M. et al. Accelerating loss of seagrasses across the globe threatens coastal ecosystems. Proc. Natl. Acad. Sci. 106, 12377–12381 (2009).

13. Macreadie, P. I. et al. Can we manage coastal ecosystems to sequester more blue carbon? Front. Ecol. Environ. 15, 206–213 (2017).

14. Pessarrodona, A. et al. Carbon sequestration and climate change mitigation using macroalgae: a state of knowledge review. Biol. Rev. brv.12990 (2023) doi:10.1111/brv.12990.

15. Vanderklift, M. A. et al. A Guide to International Climate Mitigation Policy and Finance Frameworks Relevant to the Protection and Restoration of Blue Carbon Ecosystems. Front. Mar. Sci. 9, 872064 (2022).

16. Howard, J. et al. Blue carbon pathways for climate mitigation: Known, emerging and unlikely. Mar. Policy 156, 105788 (2023).

17. Krause-Jensen, D. & Duarte, C. M. Substantial role of macroalgae in marine carbon sequestration. Nat. Geosci. 9, 737–742 (2016).

18. Duarte, C. M. et al. Global estimates of the extent and production of macroalgal forests. Glob. Ecol. Biogeogr. 31, 1422–1439 (2022).

19. Jayathilake, D. R. M. & Costello, M. J. A modelled global distribution of the kelp biome. Biol. Conserv. 252, 108815 (2020).

20. Pedersen, M. F. et al. Detrital carbon production and export in high latitude kelp forests. Oecologia 192, 227–239 (2020).

21. Ortega, A. et al. Important contribution of macroalgae to oceanic carbon sequestration. Nat. Geosci. 12, 748–754 (2019).

22. Filbee-Dexter, K., Wernberg, T., Norderhaug, K. M., Ramirez-Llodra, E. & Pedersen, M. F. Movement of pulsed resource subsidies from kelp forests to deep fjords. Oecologia 187, 291–304 (2018).

23. Queirós, A. M. et al. Identifying and protecting macroalgae detritus sinks toward climate change mitigation. Ecol. Appl. 33, e2798 (2023).

24. Krumhansl, K. & Scheibling, R. Production and fate of kelp detritus. Mar. Ecol. Prog. Ser. 467, 281– 302 (2012).

25. Hurd, C. L. et al. Forensic carbon accounting: Assessing the role of seaweeds for carbon sequestration. J. Phycol. 58, 347–363 (2022).

26. Paine, E. R., Schmid, M., Boyd, P. W., Diaz-Pulido, G. & Hurd, C. L. Rate and fate of dissolved organic carbon release by seaweeds: a missing link in the coastal ocean carbon cycle. J. Phycol. 57, 1375– 1391 (2021).

27. Geraldi, N. R. et al. Fingerprinting blue carbon: rationale and tools to determine the source of organic carbon in marine depositional environments. Front. Mar. Sci. 263 (2019).

28. Krause-Jensen, D. et al. Sequestration of macroalgal carbon: the elephant in the Blue Carbon room. Biol. Lett. 14, 20180236 (2018).

29. Queirós, A. M. et al. Connected macroalgal-sediment systems: blue carbon and food webs in the deep coastal ocean. Ecol. Monogr. 89, (2019).

30. Fujita, R. et al. Seaweed blue carbon: Ready? Or Not? Mar. Policy 155, 105747 (2023).

31. Araya-Lopez, R., de Paula Costa, M. D., Wartman, M. & Macreadie, P. I. Trends in the application of remote sensing in blue carbon science. Ecol. Evol. 13, e10559 (2023).

32. Mora-Soto, A. et al. Environmental characteristics and floating kelp dynamics in the southern Salish Sea, British Columbia, Canada. Front. Mar. Sci. (in review).

33. Archambault, P. et al. From sea to sea: Canada’s three oceans of biodiversity. PLoS One 5, e12182 (2010).

34. Ross, F. W. et al. A preliminary estimate of the contribution of coastal blue carbon to climate change mitigation in New Zealand. N. Z. J. Mar. Freshw. Res. 1–11 (2023).

35. Filbee-Dexter, K. & Scheibling, R. E. Sea urchin barrens as alternative stable states of collapsed kelp ecosystems. Mar. Ecol. Prog. Ser. 495, 1–25 (2014).

36. Thompson, M. MaPP Kelp Monitoring Protocol. (2021).

37. Filbee-Dexter, K. et al. Ocean temperature controls kelp decomposition and carbon sink potential. PLOS Biol. (2020).

38. Filbee-Dexter, K. et al. Carbon export from seaweed forests to deep ocean sinks. (in review).

39. Connell, S. D. et al. Recovering a lost baseline: missing kelp forests from a metropolitan coast. Mar. Ecol. Prog. Ser. 360, 63–72 (2008).

40. Filbee-Dexter, K., Feehan, C. & Scheibling, R. Large-scale degradation of a kelp ecosystem in an ocean warming hotspot. Mar. Ecol. Prog. Ser. 543, 141–152 (2016).

41. Johnson, C. R. et al. Climate change cascades: Shifts in oceanography, species’ ranges and subtidal marine community dynamics in eastern Tasmania. J. Exp. Mar. Biol. Ecol. 400, 17–32 (2011).

42. Steneck, R. S. et al. Kelp forest ecosystems: biodiversity, stability, resilience and future. Environ. Conserv. 29, 436–459 (2002).

43. Wernberg, T. et al. An extreme climatic event alters marine ecosystem structure in a global biodiversity hotspot. *Nat*. Clim. Change 3, 78–82 (2013).

44. Starko, S. et al. Microclimate predicts kelp forest extinction in the face of direct and indirect marine heatwave effects. Ecol. Appl. 32, (2022).

45. Steneck, R. S., Leland, A., McNaught, D. C. & Vavrinec, J. Ecosystem flips, locks, and feedbacks: the lasting effects of fisheries on Maine’s kelp forest ecosystem. Bull. Mar. Sci. 89, 31–55 (2013).

46. Attridge, C. M., Metaxas, A. & Denley, D. Wave exposure affects the persistence of kelp beds amidst outbreaks of the invasive bryozoan Membranipora membranacea. Mar. Ecol. Prog. Ser. 702, 39–56 (2022).

47. Oliver, E. C. et al. Projected marine heatwaves in the 21st century and the potential for ecological impact. Front. Mar. Sci. 6, 734 (2019).

48. Assis, J., Araújo, M. B. & Serrão, E. A. Projected climate changes threaten ancient refugia of kelp forests in the North Atlantic. Glob. Change Biol. 24, e55–e66 (2018).

49. Jueterbock, A. et al. Climate change impact on seaweed meadow distribution in the North Atlantic rocky intertidal. Ecol. Evol. 3, 1356–1373 (2013).

50. Filbee-Dexter, K. & Wernberg, T. Substantial blue carbon in overlooked Australian kelp forests. Sci. Rep. 10, 12341 (2020).

51. Goldsmit, J. et al. Kelp in the Eastern Canadian Arctic: Current and Future Predictions of Habitat Suitability and Cover. Front. Mar. Sci. 18, 742209 (2021).

52. Wright, L., Pessarrodona, A. & Foggo, A. Climate-driven shifts in kelp forest composition reduce carbon sequestration potential. Glob. Change Biol. (2022).

53. Filbee-Dexter, K., Wernberg, T., Fredriksen, S., Norderhaug, K. M. & Pedersen, M. F. Arctic kelp forests: Diversity, resilience and future. Glob. Planet. Change 172, 1–14 (2019).

54. Starko, S. et al. Temperature and food chain length, but not latitude, explain region-specific kelp forest responses to an unprecedented heatwave. bioRxiv 2023–01 (2023).

55. Pessarrodona, A., Filbee-Dexter, K., Krumhansl, K. A., Moore, P. J. & Wernberg, T. A global dataset of seaweed net primary productivity. Sci. Data (2022) doi:10.1101/2021.07.12.452112.

56. Filbee-Dexter, K., et al. *in review.* Seaweed forests are carbon sinks that can mitigate CO2 emissions. Nature Geosciences.

57. Leithold, E. L., Blair, N. E. & Wegmann, K. W. Source-to-sink sedimentary systems and global carbon burial: A river runs through it. Earth-Sci. Rev. 153, 30–42 (2016).

58. Hurd, C., Gattuso, J. & Boyd, P. W. Air-sea carbon dioxide equilibrium: Will it be possible to use seaweeds for carbon removal offsets? J. Phycol. (2023).

59. Needelman, B. A. et al. The science and policy of the verified carbon standard methodology for tidal wetland and seagrass restoration. Estuaries Coasts 41, 2159–2171 (2018).

60. Starko, S. et al. A comprehensive kelp phylogeny sheds light on the evolution of an ecosystem. Mol. Phylogenet. Evol. 136, 138–150 (2019).

61. Druehl, L. D. The pattern of Laminariales distribution in the northeast Pacific. Phycologia 9, 237–247 (1970).

62. Bringloe, T. T., Verbruggen, H. & Saunders, G. W. Population structure in Arctic marine forests is shaped by diverse recolonisation pathways and far northern glacial refugia. bioRxiv 2020–03 (2020).

63. Bolton, J. J. The biogeography of kelps (Laminariales, Phaeophyceae): a global analysis with new insights from recent advances in molecular phylogenetics. Helgol. Mar. Res. 64, 263–279 (2010).

64. Assis, J. et al. A fine-tuned global distribution dataset of marine forests. Sci. Data 7, 119 (2020).

65. Pessarrodona, A. et al. Global seaweed productivity. Sci. Adv. 8, eabn2465 (2022).

66. Castro de la Guardia, L., et al. Increasing depth distribution of Arctic kelp with increasing number of open water days with light. Elem Sci Anth 11, 00051 (2023).

67. Greenlaw, M. & Harvey, C. Data of: A substrate classification for the Inshore Scotian Shelf and Bay of Fundy, Maritimes Region. (2022).

68. Gregr, E. J., Haggarty, D. R., Davies, S. C., Fields, C. & Lessard, J. Comprehensive marine substrate classification applied to Canada’s Pacific shelf. PLOS ONE 16, e0259156 (2021).

69. Assis, J. et al. Bio-ORACLE v2. 0: Extending marine data layers for bioclimatic modelling. Glob. Ecol. Biogeogr. 27, (2018).

70. Dive Survey Algae And Substrate Data. OpenData Fisheries & Oceans Canada (2024).

71. Krumhansel, K. Unpublished data. (2022).

72. British Columbia Shore Zone Data Portal. https://soggy2.zoology.ubc.ca/geonetwork/srv/eng/catalog.search#/metadata/501b77d2-573c-44d9-ac92-140d0f024e44 (2022).

73. Howes, D., Harper, J. & Owens, E. Physical shore-zone mapping system for British Columbia. Rep. Prep. Environ. Emerg. Serv. Minist. Environ. Vic. BC Coast. Ocean Resour. IncSidney BC Owens Coast. Consult. Bainbridge WA (1994).

74. Mora-Soto, A. et al. A high-resolution global map of giant kelp (*Macrocystis pyrifera*) forests and intertidal green algae (*Ulvophyceae*) with Sentinel-2 imagery. Remote Sens. 12, 694 (2020).

75. Bürkner, P.-C. Advanced Bayesian multilevel modeling with the R package brms. ArXiv Prepr.ArXi*v170511123* (2017).

76. Bürkner, P.-C. brms: An R package for Bayesian multilevel models using Stan. J. Stat. Softw. 80, 1–28 (2017).

77. Barreda, S. & Silbert, N. Fitting Bayesian regression models with brms. in Bayesian Multilevel Models for Repeated Measures Data 55–86 (Routledge, 2023).

78. Gelman, A. Parameterization and Bayesian modeling. J. Am. Stat. Assoc. 99, 537–545 (2004).

79. Vehtari, A., Gelman, A. & Gabry, J. Practical Bayesian model evaluation using leave-one-out cross-validation and WAIC. Stat. Comput. 27, 1413–1432 (2017).

80. Vehtari, A., et al. loo: Efficient leave-one-out cross-validation and WAIC for Bayesian models. (2023).

