## Appendix A for "A blueprint for national assessments of the blue carbon capacity of kelp forests applied to Canada’s coastline"

We present a blueprint for conducting a national assessment of the carbon sequestration capacity of kelp forest ecosystems (Figure A1). Recognizing the significant data gaps and uncertainties that all countries will face in this regard, our reproductible and transparent blueprint workflow involves four main stages. In the first stage, the aim is to extensively collate and synthesize available kelp datasets to identify key data sources and to reveal any data gaps that may hinder the reliable estimation of kelp carbon sequestration processes on a national scale. In the second stage, the aim is to evaluate the potential for natural variation in per-area carbon stocks and fluxes (e.g., productivity) across kelp species and different oceanographic contexts. In the third stage, the aim is to produce an initial estimate of the total carbon stock, production, and sequestration capacity of kelp forests at a national scale, acknowledging the range of potential uncertainties involved. The final stage is to fill current data gaps through targeted sampling and to re-evaluate estimates as new information becomes available. Below is a step-by-step guide, outlining each stage of the assessment, with special considerations and resources for replicating the process in data-limited kelp-dominated systems.

Stage:1 Compile and synthesize available data

- **An important first step is to define the scope of the assessment, including the ecological focus and study domain.** Researchers need to decide what kelp species to include in the assessment, which will ultimately determine the extent of the study domain. Certain countries may primarily have subsurface kelps, while countries on the western boundaries of continents could have surface kelps in addition to subsurface kelp forests, extending from the subtidal into the intertidal zone. National monitoring programs and global kelp occurrence databases can help to identify the primary kelp species found within national jurisdictions[1,2]. Web portals, such as Algaebase (https://algaebase.org) can also aid this process. An appropriate study domain for subtidal kelp forests could extend from mean-low-water out to a specified depth that is informed by *in situ* observations and data describing the maximum depth limit of kelps. Where there is appropriate data, researchers may also choose to extend the study domain to the mean-high-water mark to capture the potential role of intertidal kelps.
- **The next step is to collate available information and datasets on the biomass, plant density, canopy cover, and net primary productivity (NPP) of kelp species from published and unpublished sources**. This step is meant to identify key datasets available for the assessment and to reveal any important data gaps that must be filled to reliably estimate the carbon stocks and production rates associated with kelp species. All NPP (kg DW or WW $m^{-2}$ $Yr^{-1}$) and abundance data (e.g., kg DW or WW $m^{-2}$) can be converted

to carbon stock (Mg C $Ha^{-1}$) and annual carbon production (Mg C $Ha^{-1}$ $Yr^{-1}$) estimates using published species-specific and coast-specific conversions (i.e., wet-dry ratios and carbon-dry weight ratios) from the peer reviewed literature.[3] Plant density and canopy cover datasets are also needed to develop biologically constrained estimates of kelp forest extents, stocks, and production rates across coasts. Researchers can use an existing macroalgal NPP database[3] to obtain available information on the NPP rates of kelp species and to structure a search of the peer reviewed literature for other necessary kelp datasets. Existing kelp forest monitoring program data and unpublished datasets can also be useful sources of kelp abundance datasets (i.e., biomass, density, and percent cover).

- **The next step is to use available information and data to estimate the total areal extent of kelp forests.** Areal extent estimates will be needed to extrapolate estimates of kelp carbon stocks, carbon production rates, and carbon export rates to a national scale. For surface kelps, areal extent can be obtained using available high resolution remote sensing products and species distribution modeling studies[1,2]. In the absence of such data, a suitable reef approach can be employed to obtain a high-, mid-, and low-end estimate for potential extent of kelp forests. Calculating the area of available rocky reef habitat that is above the maximum depth limit of kelp species can provide a useful estimate of the hypothetical maximum limit for kelp forest distributions. However, these estimates can and should be modified using available kelp percent cover and density data to provide a more biologically constrained mid and low-end estimate. Where higher resolution bathymetry data is not available, researchers can obtain grided depth data from the General Bathymetric Chart of the Ocean database (https://www.gebco.net/). Additionally, where there is a lack of grided substrate data within national jurisdictions, coarse proxies can be obtained from global datasets[4,5].
- **A final step is to compile any available data can allow for the approximation of kelp carbon fluxes to and sequestration rates in various potential sinks.** Currently there is limited data on the potential kelp carbon fluxes and accumulation of kelp derived POC and DOC in shelf sediments and the deep ocean. Coarse global estimates can be obtained from Krause-Jensen et al. 2016[6], although with considerable uncertainty. More resolved estimates of the rate of carbon export (% of kelp production) to the continental shelf break (i.e., below 200 m depth) are also available from ocean transport models of shelf to open ocean exchange (e.g., [7]), which can serve to approximate the amount of kelp carbon reaching the deep ocean at an ecoregional scale. However, estimates from ocean transport models still require a thorough interrogation with in-situ experiments and observation studies.

Stage 2: Quantify natural variation in carbon stock, production, and export

- **An important first step here is to evaluate the potential for natural variation within and across kelp species.** We used Bayesian hierarchical models, implemented using the ‘brms’ package of R, to produce parameter estimates of the average carbon stocks and production rates of different kelp species. Bayesian hierarchical models are parameterized similarly to hierarchical linear regression models. The observed carbon stocks and production rates of different kelp species are treated as the response variables, while environmental variables (e.g., sea surface temperature, ocean, ecoregion) that potentially influence rates of short-term biomass accumulation and primary productivity are included as fixed effects. Site- and study-level variation can also be accommodated through a model hierarchical structure (i.e., as random effects). A key advantage of using Bayesian approaches is that they provide a posterior distribution a range of possible estimated values given prior beliefs and available data. These prior beliefs can be informed by observations from different systems and species. From this posterior distribution, researchers can calculate a posterior mean and a credible confidence interval (CCI) indicating the range within which the true parameter value is most likely to fall. Additionally, Bayesian models can consider measurements of standard deviation as an additional response term, allowing for adjusted CCIs that reflect greater uncertainty where there is greater variability in kelp biomass and NPP. Researchers can use these credible confidence intervals to identify significant differences in carbon stocks and production rates across kelp species and coasts. Additional information about using the R package "brms" can be found in the literature and the documentation[8,9]. Scripts for model parameterization, selecting informative priors, and evaluating model outputs, can also found in our Github repository.

- **The next step is to evaluate the potential for natural variation within and across different regional contexts.** To account for the fact the kelp forests are often multi-species assemblages with different relative compositions, we recommend determining the per-area carbon stocks and production rates of kelp forests as the summed posterior mean estimates for the kelp species found in a particular location, weighted by their observed relative abundance. Additionally, per-area carbon fluxes from kelp forests can be approximated using data on the fraction of kelp detritus that is likely to reach the continental shelf break (i.e., 200 m) from open ocean exchange models. While export does not directly equate to carbon sequestration it can provide a useful proxy in the absence of direct measurements of kelp carbon sequestration in the deep ocean. Where there is limited information on the occurrence and composition of kelp forest communities, a first order estimate could be calculated at the coast scale (e.g., for the Pacific coast of British Columbia). However, where there are more comprehensive datasets, it is recommended to produce more spatially resolved estimates (e.g., at the ecoregion scale).

Stage 3: Develop initial national estimates of the carbon sequestration capacity of kelp forests

- **Using the outputs of stages 1 and 2, the next step is to produce estimates of the carbon sequestration capacity of kelp forests at a national scale.** The total standing carbon stock, production, and export capacity of kelp forests can be determined by multiplying the per-area estimates by the areal extent estimates. Since surface and subsurface kelps may co-occur, we recommend treating these groups separately. In data limited systems, a first order estimate would be to produce estimates at the coast scale, but in more data rich systems it may be possible to produce estimates at the ecoregion scale. Then national estimates can be determined by summing across coasts and ecoregions.

- **An important step is to quantify the sensitivity and range of uncertainties surrounding the national estimates.** Given the current data limitations and difficulties with estimating kelp carbon sequestration processes, assessors may need to use a set of reasonable assumptions. For example, in systems lacking comprehensive habitat maps, assessors may need to make assumptions about the maximum depth limit of kelps, the prevalence of rocky reefs, and the relative abundance of kelp species when estimating the areal extent of subsurface kelp forests. Conducting a sensitivity analysis, where the parameters used are systematically varied, can help evaluate the impact of these choices on the results and the inherent uncertainties involved. Sensitivity analyses can also help identify the highest priorities for future data collection.

Stage 4: Re-evaluate estimates based on new information and data

- **Re-evaluation is integral to ensuring that the assessment remains relevant and accurate over time**. Advances in technology, improved monitoring, and additional research are likely to enhance our understanding of kelp carbon sequestration processes. Researchers should assess the impact of any changes in methodology, data availability, and environmental conditions on the resulting estimates.
- **Collaboration and transparency are crucial to the revaluation process.** Engaging with experts and incorporating diverse perspective ensures that the assessment benefits from a comprehensive understanding of kelp ecosystems. Clearly documenting any updates, modifications, or changes made to the methodology will ensure the reproductivity and reliability of the assessment. Open communication about the uncertainties and limitations associated with the estimates will help to build trust in the scientific process and the resulting findings.

Conclusion

This blueprint offers a reproducible and transparent guide for assessing the carbon capacity of kelp forests in national jurisdictions. Researchers can adipate these methods, considering the regional context and data gaps, to contribute valuable insights into the capacity for kelp forests to support NCS at national scales. The four-stage workflow outline in this blueprint aims to address challenges posed by data limitations and uncertainties, promoting a rigorous and iterative approach to enhance the reliability of estimates over time.

**Figure A1.** A proposed blueprint for national assessments of the blue carbon potential of kelp forests.

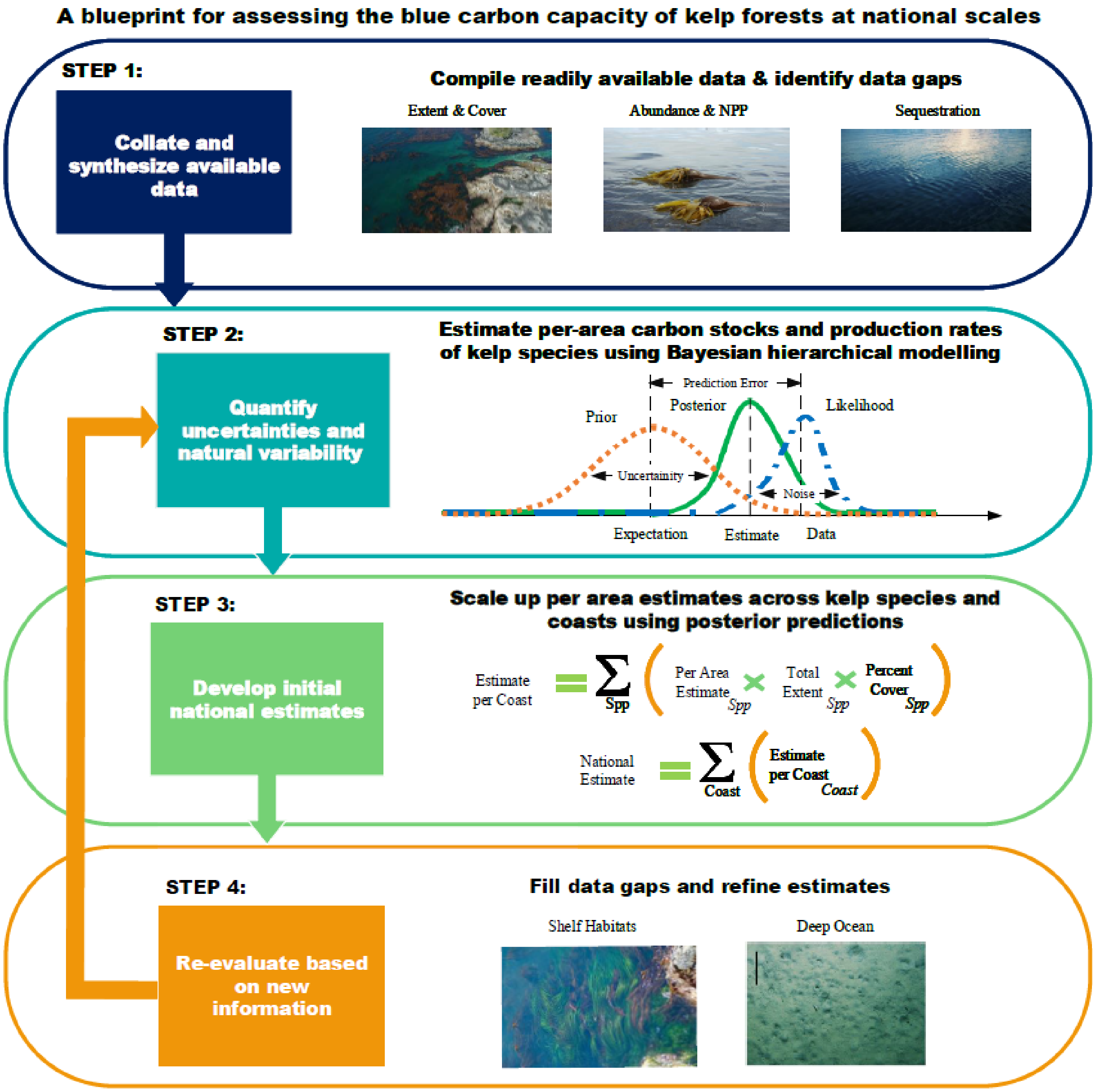

**References:**


1. Jayathilake, D. R. M. & Costello, M. J. A modelled global distribution of the kelp biome. *Biological Conservation* **252**, 108815 (2020).
2. Assis, J. *et al.* A fine-tuned global distribution dataset of marine forests. *Sci Data* **7**, 119 (2020).
3. Pessarrodona, A. *et al.* Global seaweed productivity. *Sci. Adv.* **8**, eabn2465 (2022).
4. Young, A. P. & Carilli, J. E. Global distribution of coastal cliffs. *Earth Surface Processes and Landforms* **44**, 1309–1316 (2019).
5. Duarte, C. M. *et al.* Global estimates of the extent and production of macroalgal forests. *Global Ecol. Biogeogr.* **31**, 1422–1439 (2022).
6. Krause-Jensen, D. & Duarte, C. M. Substantial role of macroalgae in marine carbon sequestration. *Nature Geosci* **9**, 737–742 (2016).
7. Filbee-Dexter, K. *et al.* Carbon export from seaweed forests to deep ocean sinks. (in review).
8. Bürkner, P.-C. brms: An R package for Bayesian multilevel models using Stan. *Journal of statistical software* **80**, 1–28 (2017).
9. Barreda, S. & Silbert, N. Fitting Bayesian regression models with brms. in *Bayesian Multilevel Models for Repeated Measures Data* 55–86 (Routledge, 2023).
