## Appendix C for "A blueprint for national assessments of the blue carbon capacity of kelp forests applied to Canada’s coastline"

**Supplemental Information**

**Appendix C**

**Figures**

**Figure C1.** Distribution of datasets used in the assessment across the Pacific (purple), Arctic (blue), and Atlantic coasts (green) of Canada and the northwestern hemisphere. Kelp biomass and density records are denoted by pink circles; net primary productivity records are denoted by yellow circles; and kelp percent cover records are denoted by orange circles.

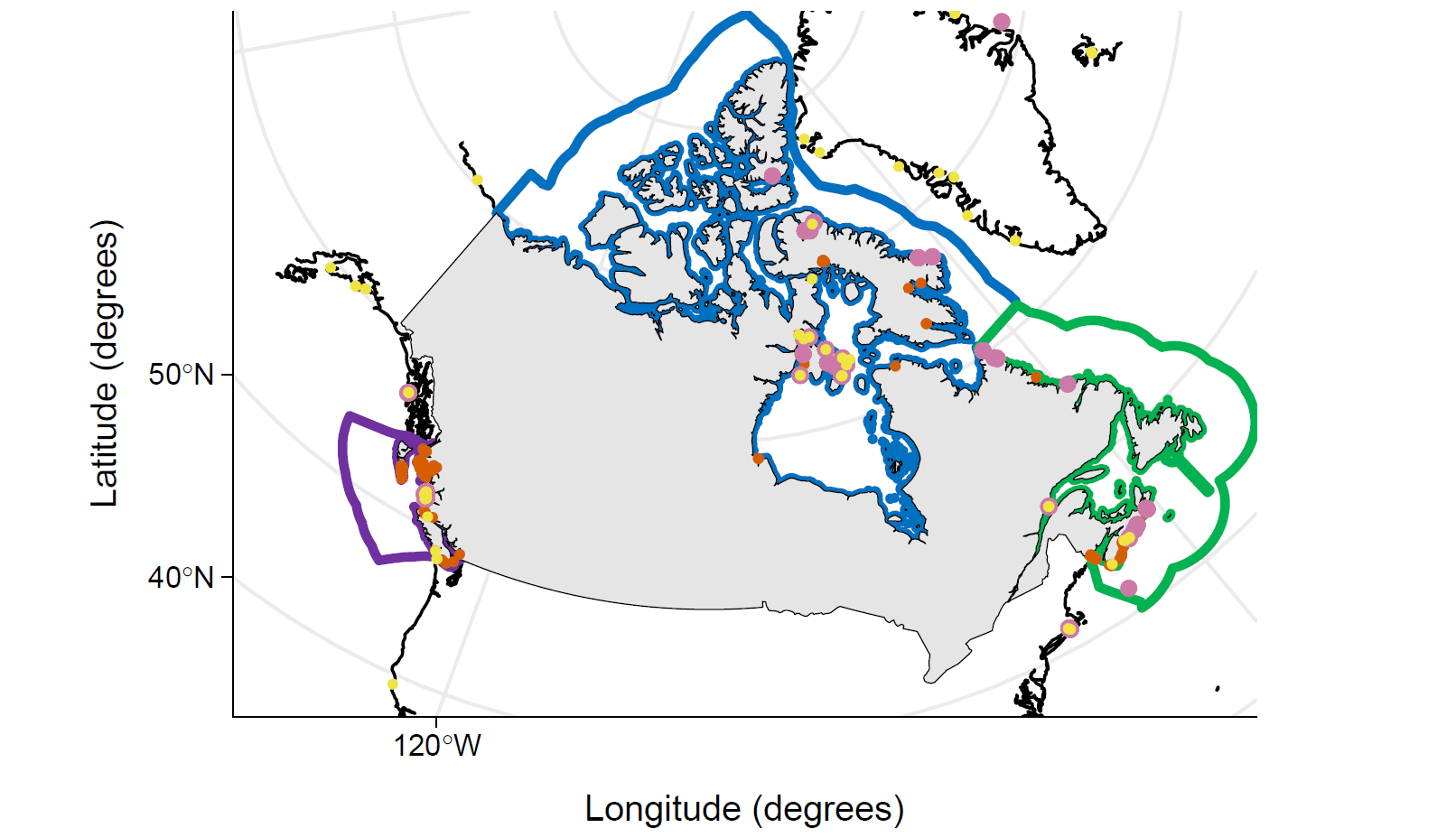

**Figure C2.** Surface kelp forest distribution according to available datasets from the Pacific Coast of Canada. a) A mid bound extent for surface kelps was determined using shoreline maps derived from aerial surveys conducted by the British Columbia ShoreZone Survey Program completed from 2004 to 2007, representing the linear shoreline extent of surface kelps. The mid bound extent was calculated as the areal intersection between the observed shoreline extent and the area of adjacent rocky reef shallower than 10m water depth. b) A low bound extent for surface kelps was determined using maps derived from remote sensing data from the Sentinel-2 program, representing the extent of surface kelps at peak biomass between 2015 and 2019.

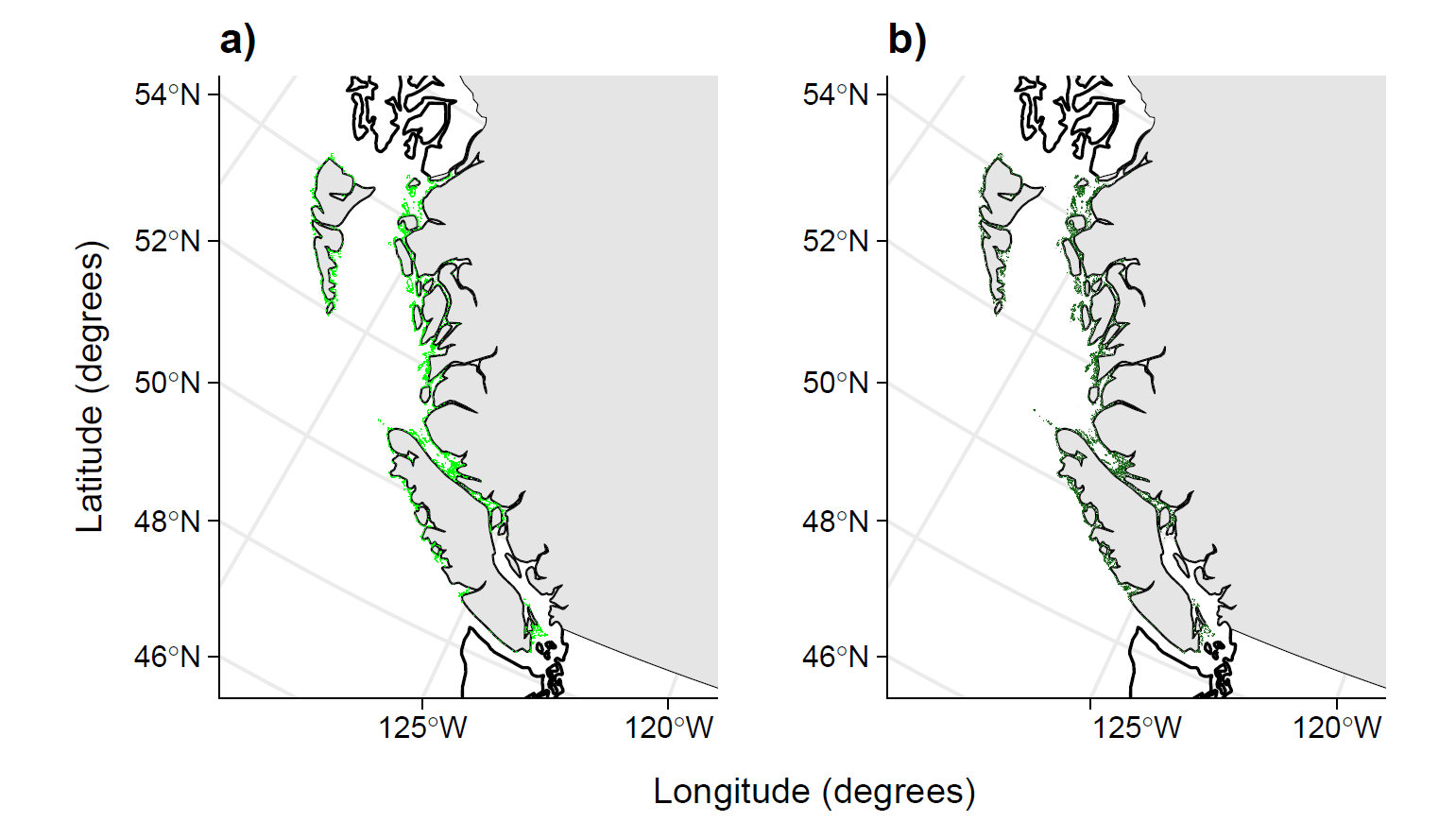

**Figure C3.** Depth distribution of surface kelps, *Macrocystis pyrifera* and *Nereocystis leutkeana,* from in-situ remotely operated vehicle surveys along the Pacific Coast of Canada. Red dots show the mean observed depth distribution for each species, with two standard deviations surrounding the mean (red whiskers).

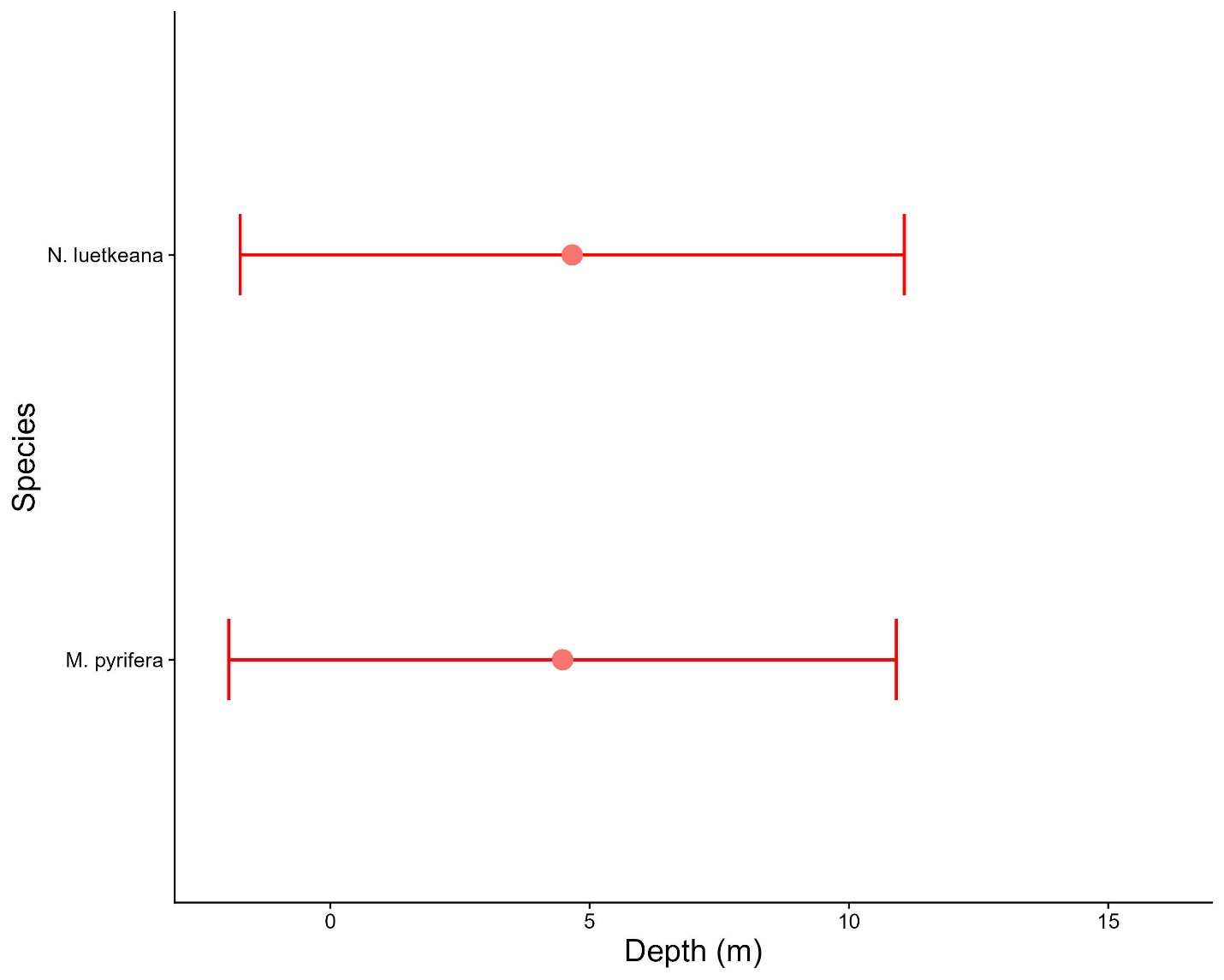

**Figure C4.** Sensitivity of areal extent estimates for subsurface kelp forests to changes in rock cover (%), kelp percent cover (%), and kelp maximum depth (m) parameters.

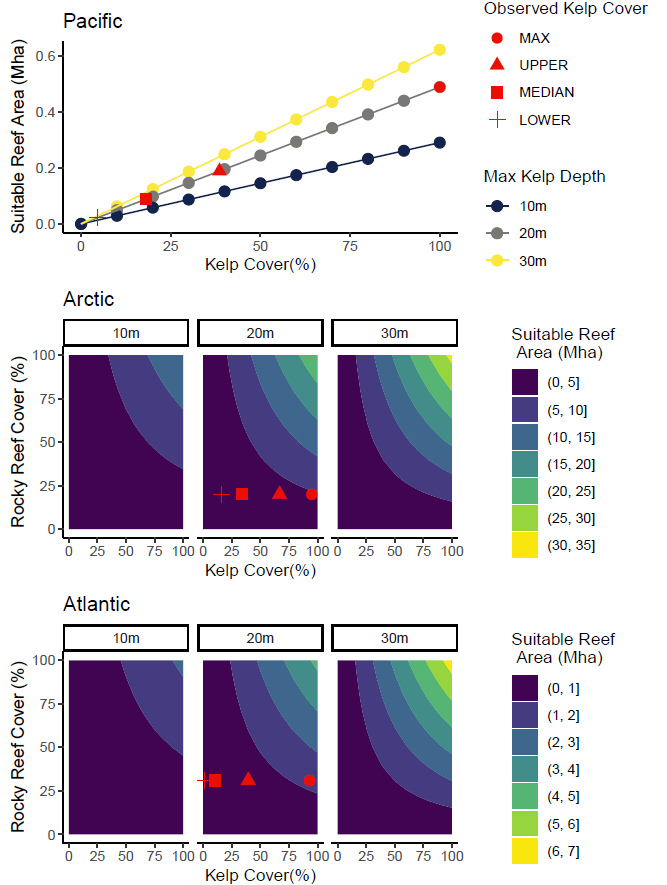

**Figure C5.** Sensitivity of standing annual carbon production estimates for subsurface kelp forests to changes in kelp area (Mha) and per area rates of kelp primary production Mg C ha^-1^ yr^-1^. Red symbols indicate the estimates representing the lower (+), median (□), upper (Δ), and maximum (○) extent of subsurface kelps in Canada.

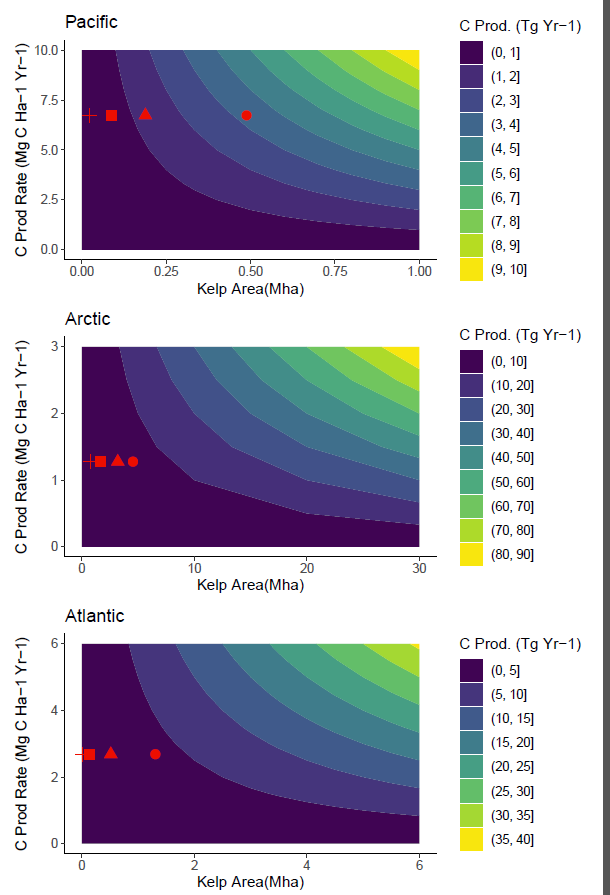

**Figure C6.** Sensitivity of standing annual carbon export estimates for subsurface kelp forests to changes in kelp area (Mha) and per area rates of kelp primary production Mg C ha^-1^ yr^-1^. Red symbols indicate the estimates representing the lower (+), median (□), upper (Δ), and maximum (○) extent of subsurface kelps in Canada.

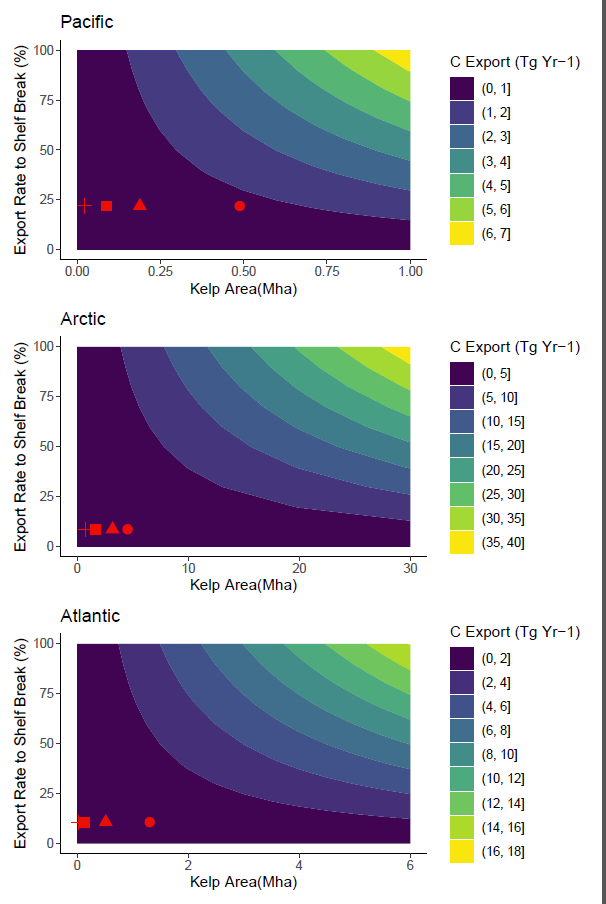

**Figure C7.** Distribution of observed (y) carbon stocks (Mg C Ha^-1^) of eleven kelp species compared to simulated data from the posterior predictive distribution (yrep) of the final Bayesian hierarchical models. Kelp species include: a) *Macrocystis pyrifera*, b), *Nereocystis leutkeana* c),*Costaria costata,* d) *Agarum clathratum / Neoagarum fimbriatum*, e) *Laminaria digitata / Hedophyllum nigripes,* f) *Laminaria solidungula*, g) *Pterygophera californica*, h) *Pleurophycus gardneri,* and i) *Saccharina latissima*.

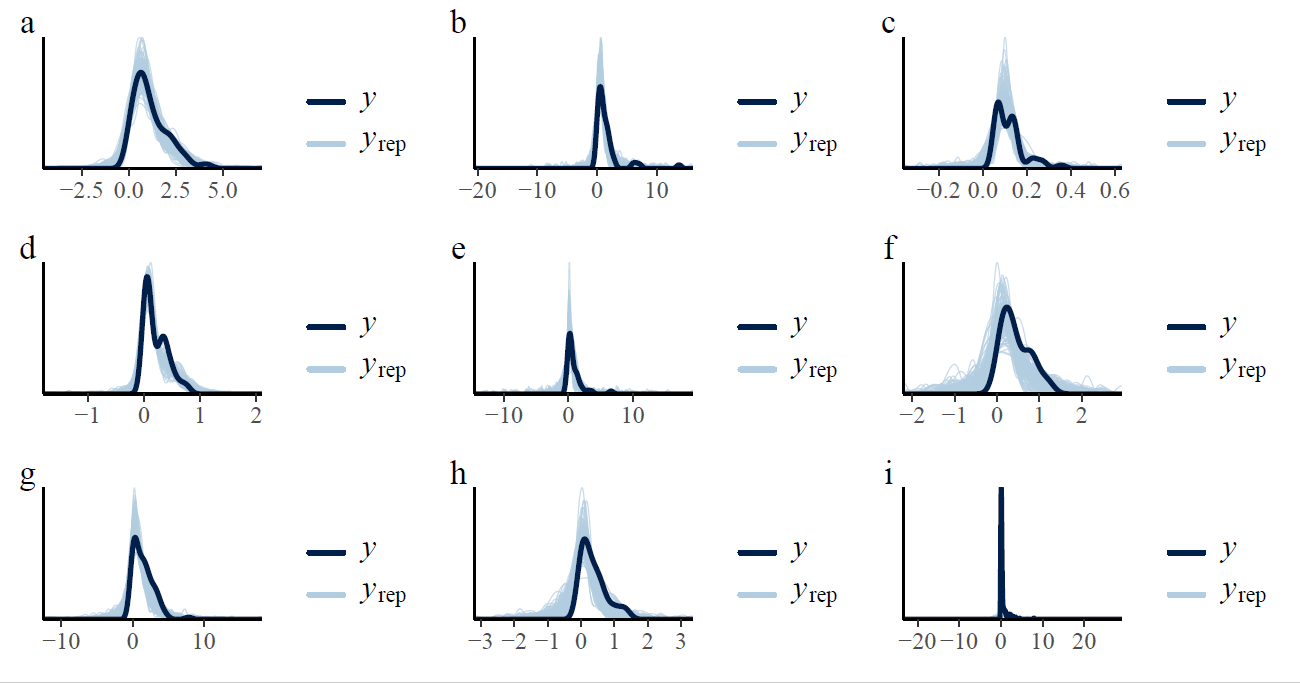

**Figure C8.** Distribution of observed (y) carbon production rates (Mg C Ha^-1^ Yr^-1^) of eleven kelp species compared to simulated data from the posterior predictive distributiona (yrep) of the final Bayesian hierarchical models. Kelp species include: a) *Macrocystis pyrifera*, b), *Nereocystis leutkeana* c),*Costaria costata,* d) *Agarum clathratum*, e) *Laminaria digitata / Hedophyllum nigripes,* f) *Laminaria solidungula*, g) *Pterygophera californica*, h) *Pleurophycus gardneri,* and i) *Saccharina latissima*.

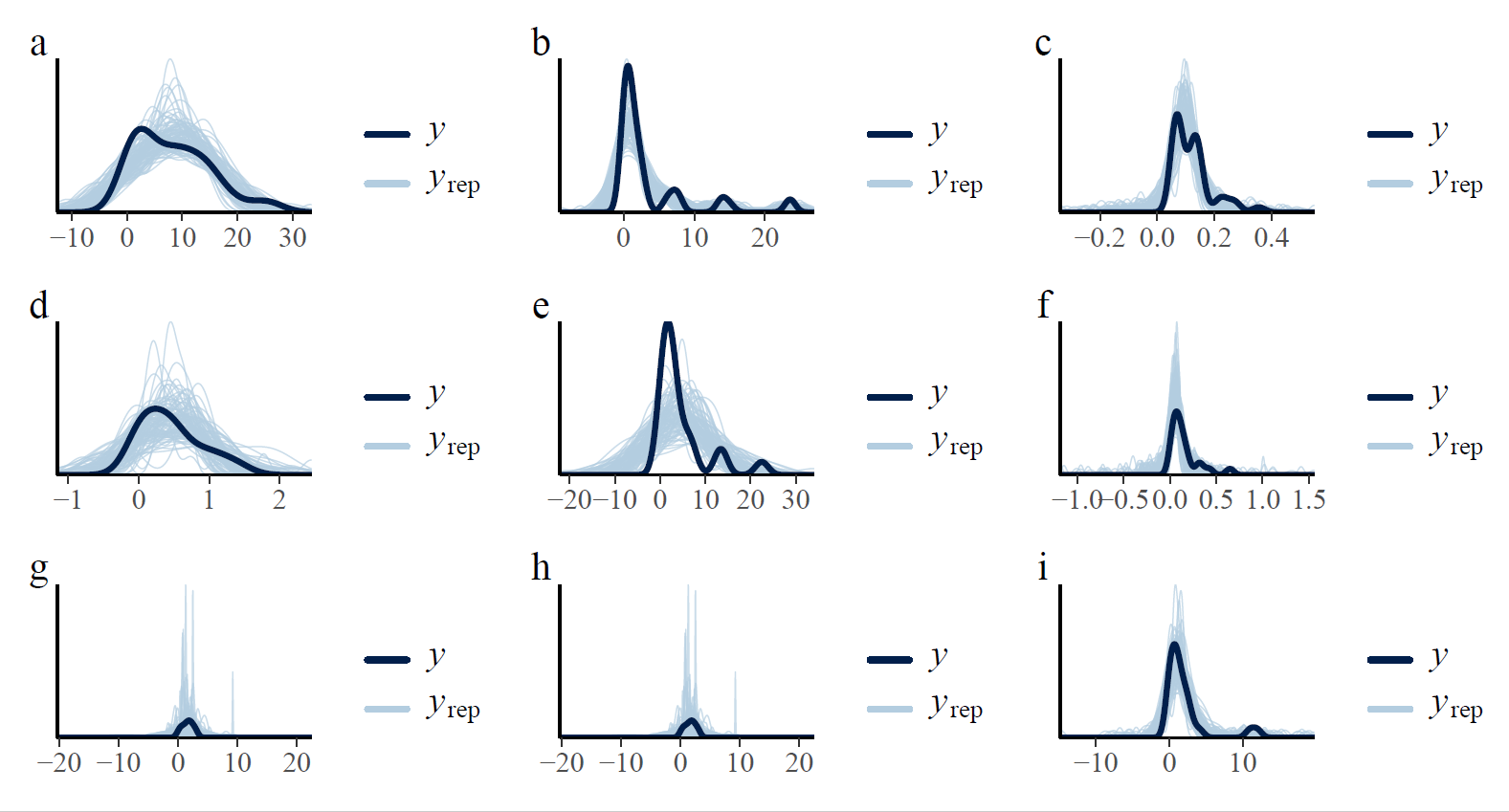

**Tables**

**Table C1.** Full summary table of data types and sources to used to identify data gaps across regions and species.

| **Scientific name** | **Aerial Extent** | **Canopy Cover** | **Canopy Biomass** | **Net Primary Production** |
| --- | --- | --- | --- | --- |
| Agarum clathratum | ^This study^ | ^1,2^ | ^2–7^ | ^4,8^ |
| Alaria esculenta |  | ^2^ | ^2^ |  |
| Costaria costata |  |  | ^6^ | ^6^ |
| Cymathaere triplicata |  |  | ^6^ | ^6^ |
| Egregia menziesii |  |  |  |  |
| Eisenia arborea |  |  | ^7,9^ |  |
| Hedophyllum nigripes | ^This study^ | ^2,10^ | ^2,4,11^ | ^11^ |
| Laminaria digitata | ^This study^ | ^1–3,10^ | ^3–5,12^ | ^12–15^ |
| Laminaria longipes |  |  |  |  |
| Laminaria setchellii |  |  | ^6^ |  |
| Laminaria solidungula | ^This study^ | ^2,10^ | ^2^ | ^16–20^ |
| Laminaria longipes |  |  |  |  |
| Macrocystis pyrifera | ^21,22^ |  | ^7,11,23,24^ | ^11,25–29^ |
| Neoagarum fimbriatum |  |  | ^11^ | ^11^ |
| Nereocystis luetkeana | ^21,22^ |  | ^4,24,30,31^ | ^4,24,30–32^ |
| Pleurophycus gardneri | ^This study^ |  | ^4,6^ | ^4^ |
| Pterygophora californica | ^This study^ |  | ^6^ | ^33^ |
| Saccharina latissima | ^This study^ | ^1–3,10^ | ^2–5,7,12,34–36^ | ^4,13,20,34,35,37^ |

**Table References:**

1.	Attridge, C. M., Metaxas, A. & Denley, D. Wave exposure affects the persistence of kelp beds amidst outbreaks of the invasive bryozoan Membranipora membranacea. *Mar. Ecol. Prog. Ser.* **702**, 39–56 (2022).

2.	Filbee-Dexter, K. *et al.* Sea Ice and Substratum Shape Extensive Kelp Forests in the Canadian Arctic. *Front. Mar. Sci.* **9**, 754074 (2022).

3.	Krumhansel, K. Unpublished data. (2022).

4.	Lees, D. C., Houghton, J. P., Erickson, D. E., Driskell, W. B. & Boettcher, D. E. *Ecological Studies of Intertidal and Shallow Subtidal Habitats in Lower Cook Inlet, Alaska*. https://www.osti.gov/biblio/6158236 (1980).

5.	MacGregor, K. & Johnson, L. Unpublished data. (2021).

6.	Pontier, O., Burt, J., Krumhansel, K., Okamoto, D. K. & Hessing-Lewis, M. Understory kelp biomass data from BC Central Coast v1.2.0. (2023).

7.	Yakimishyn, J. Kelp Density Monitoring - Pacific Rim National Park Reserve. (2018).

8.	Cottier, D., MacGregor, K., McKindsey, C. W. & Johnson, L. Unpublished data. (2022).

9.	Kamohara, S. *et al.* Annual net production and annual carbon and nitrogen absorptions of Eisenia arborea in the eastern coast of Ise Bay. *Nippon Suisan Gakkaishi* **75**, 1027–1035 (2009).

10.	Goldsmit, J. *et al.* Kelp in the Eastern Canadian Arctic: Current and Future Predictions of Habitat Suitability and Cover. *Front. Mar. Sci.* **18**, 742209 (2021).

11.	Bell, L. E. & Kroeker, K. J. Standing Crop, Turnover, and Production Dynamics of Macrocystis pyrifera and Understory Species Hedophyllum nigripes and Neoagarum fimbriatum in High Latitude Giant Kelp Forests. *J. Phycol.* **58**, 773–788 (2022).

12.	Smith, B. D. Comparison of Productivity Estimates for Laminaria in Nova Scotia. *Can. J. Fish. Aquat. Sci.* **45**, 557–562 (1988).

13.	Krumhansl, K. & Scheibling, R. Production and fate of kelp detritus. *Mar. Ecol. Prog. Ser.* **467**, 281–302 (2012).

14.	Pessarrodona, A. Unpublished Data.

15.	King. Unpublished Data.

16.	Chapman, A. R. O. & Lindley, J. E. Productivity of Laminaria solidungula J. Ag. in the Canadian high Arctic: a year-round study. in *Proceedings - International Seaweed Symposium* (1980).

17.	Dunton, K. H. AN ANNUAL CARBON BUDGET FOR AN ARCTIC KELP COMMUNITY. in *The Alaskan Beaufort Sea* (eds. Barnes, P. W., Schell, D. M. & Reimnitz, E.) 311–325 (Academic Press, 1984). doi:10.1016/B978-0-12-079030-2.50021-6.

18.	Dunton, K. H., Reimnitz, E. & Schonberg, S. An Arctic Kelp Community in the Alaskan Beaufort Sea. *Arctic* **35**, 465–484 (1982).

19.	Dunton, K. H. & Schell, D. M. Seasonal carbon budget and growth of Laminaria solidungula in the Alaskan High Arctic. *Mar. Ecol. Prog. Ser.* **31**, 57–66 (1986).

20.	Filbee-Dexter, K. Unpublished Data. (2019).

21.	Howes, D., Harper, J. & Owens, E. Physical shore-zone mapping system for British Columbia. *Rep. Prep. Environ. Emerg. Serv. Minist. Environ. Vic. BC Coast. Ocean Resour. IncSidney BC Owens Coast. Consult. Bainbridge WA* (1994).

22.	Mora-Soto, A. *et al.* A high-resolution global map of giant kelp (*Macrocystis pyrifera*) forests and intertidal green algae (*Ulvophyceae*) with Sentinel-2 imagery. *Remote Sens.* **12**, 694 (2020).

23.	Pontier, O., Burt, J., Okamoto, D. K. & Hessing-Lewis, M. Macrocystis kelp canopy productivity data from BC Central Coast, v1.3.0. (2023).

24.	Sutherland, I. R., Karpouzi, V., Mamoser, M. & Carswell, B. Kelp Inventory 2007- Areas of the British Columbia Central Coast from Hakai Passage to the Bardswell Group. (2008).

25.	Gerard, V. A. Some Aspects of Material Dynamics and Energy Flow in a Kelp Forest in Monterey Bay, California. (University of California, Santa Cruz, 1976).

26.	Sanderson, J. C. Subtidal macroalgal studies in East and South Eastern Tasmanian coastal waters. (University of Tasmania, 1990).

27.	Van Tussenbroek, B. I. Plant and frond dynamics of the giant kelp, Macrocystis pyrifera, forming a fringing zone in the Falkland Islands. *Eur. J. Phycol.* **28**, 161–165 (1993).

28.	Wheeler, W. N. & Druehl, L. D. Seasonal growth and productivity of *Macrocystis integrifolia* in British Columbia, Canada. *Mar. Biol.* **90**, 181–186 (1986).

29.	Rassweiler, A., Reed, D. C., Harrer, S. L. & Nelson, J. C. Improved estimates of net primary production, growth, and standing crop of Macrocystis pyrifera in Southern California. *Ecology* **99**, 2132–2132 (2018).

30.	Foreman, R. E. Studies on *Nereocystis* growth in British Columbia, Canada. *Hydrobiologia* **116**, 325–332 (1984).

31.	Pontier, O., Burt, J., Okamoto, D. K. & Hessing-Lewis, M. Nereocystis kelp canopy productivity data from BC Central Coast, v1.2.0. (2023).

32.	Weigel, B. L. & Pfister, C. A. The dynamics and stoichiometry of dissolved organic carbon release by kelp. *Ecology* **102**, e03221 (2021).

33.	Miller, R. J., Reed, D. C. & Brzezinski, M. A. Partitioning of primary production among giant kelp (Macrocystis pyrifera), understory macroalgae, and phytoplankton on a temperate reef. *Limnol. Oceanogr.* **56**, 119–132 (2011).

34.	Borum, J., Pedersen, M., Krause-Jensen, D., Christensen, P. & Nielsen, K. Biomass, photosynthesis and growth of Laminaria saccharina in a high-arctic fjord, NE Greenland. *Mar. Biol.* **141**, 11–19 (2002).

35.	Brady-Campbell, M. M., Campbell, D. B. & Harlin, M. M. Productivity of kelp (Laminaria spp.) near the southern limit in the Northwestern Atlantic Ocean. *Mar. Ecol. Prog. Ser.* **18**, 79–88 (1984).

36.	Johnson, S. W., Murphy, Csepp, Harris & Thedinga. A survey of fish assemblages in eelgrass and kelp habitats of southeastern Alaska. *US Dep. Commer. NOAA Tech. Memo* (2003).

37.	Johnston, C. S., Jones, R. G. & Hunt, R. D. A seasonal carbon budget for a laminarian population in a Scottish sea-loch. *Helgoländer Wiss. Meeresunters.* **30**, 527–545 (1977).

**Table C2.** Summary table of kelp species with sufficient data on abundance (i.e., biomass and/or density) and net primary productivity across species and coasts to be included in the assessment. Kelp icons indicate there is at least one data record for a given species and data type for a given coast. Colors of kelp icons indicate the coast from which the data was originally collected (Pacific= purple, Arctic= blue, and Atlantic = green).

| **Scientific name** | **Abundance** | **Net Primary Productivity** |
| --- | --- | --- |
| *Agarum clathratum* | 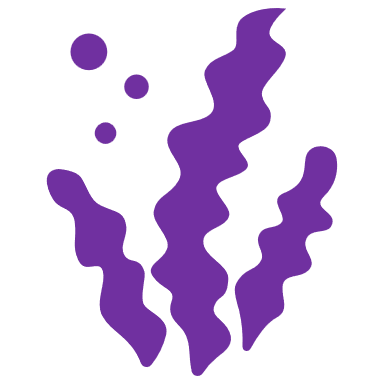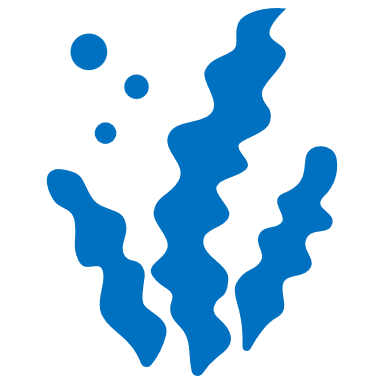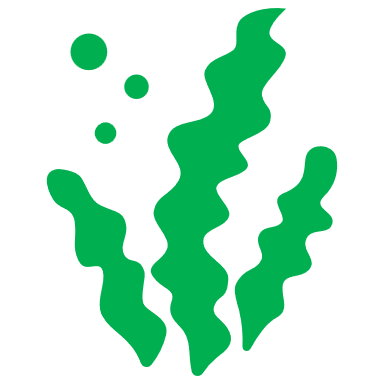 | 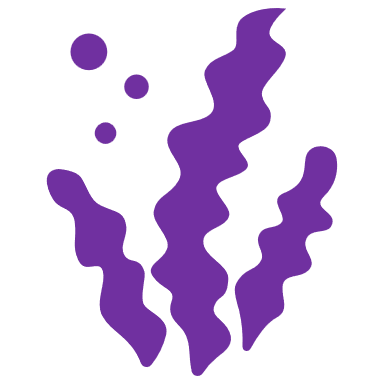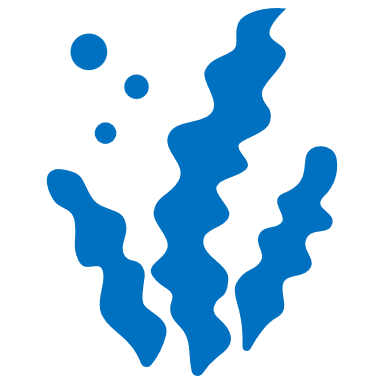 |
| *Costaria costata* | 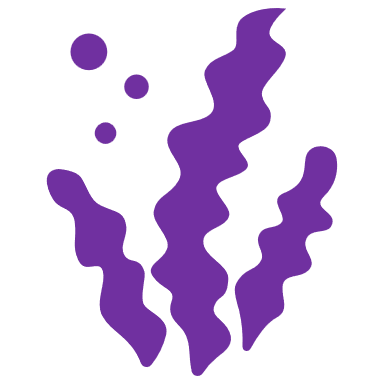 | 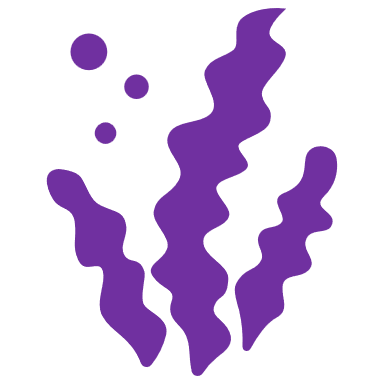 |
| *Hedophyllum nigripes* | 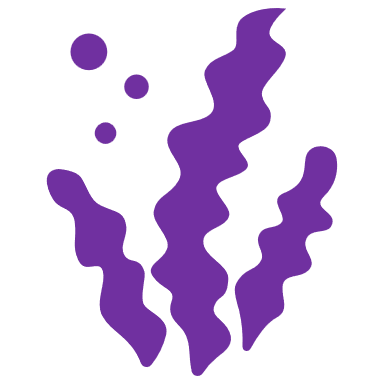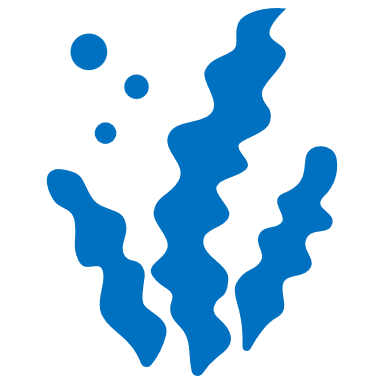 | 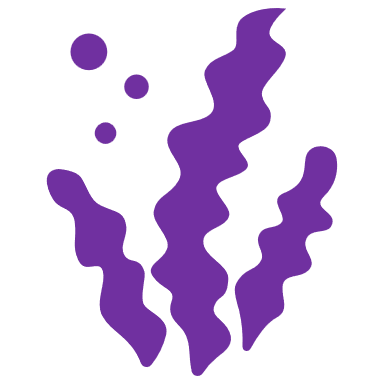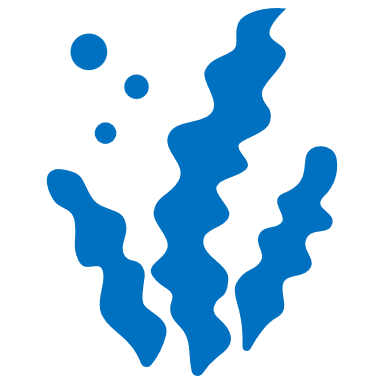 |
| *Laminaria digitata* | 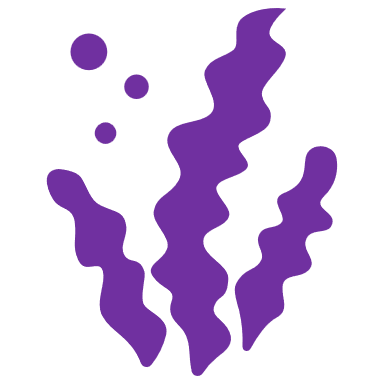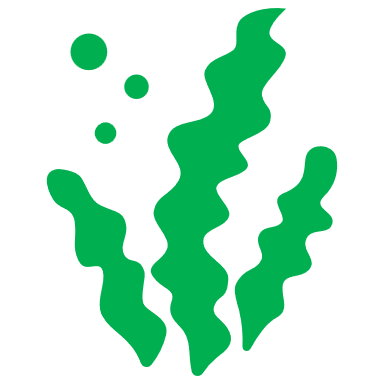 | 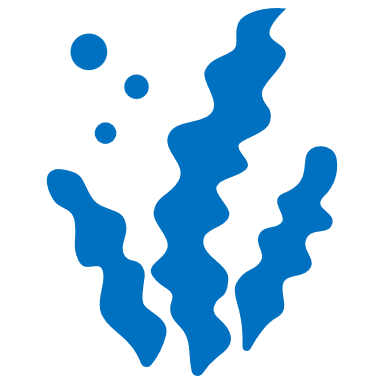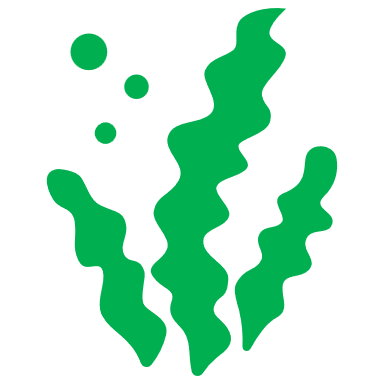 |
| *Laminaria solidungula* | 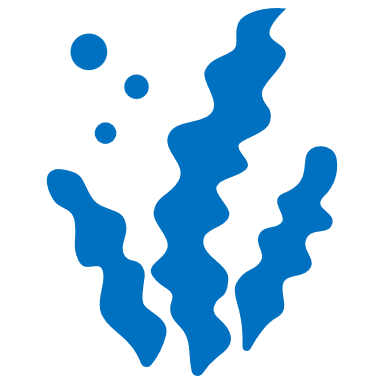 | 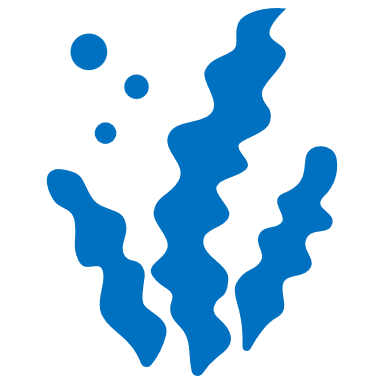 |
| *Macrocystis pyrifera* | 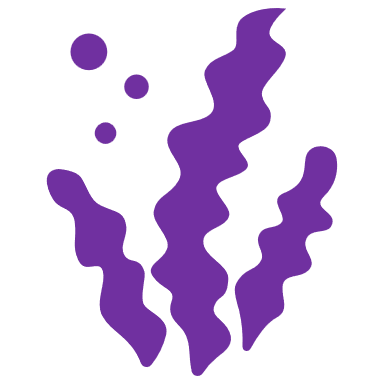 | 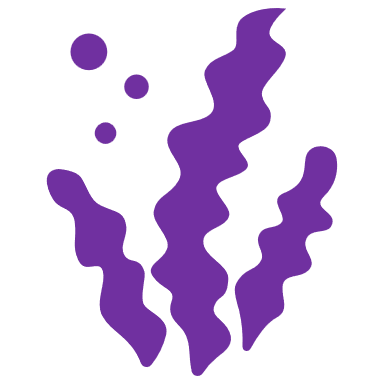 |
| *Neoagarum fimbriatum* | 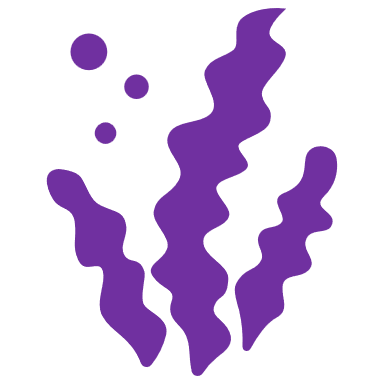 | 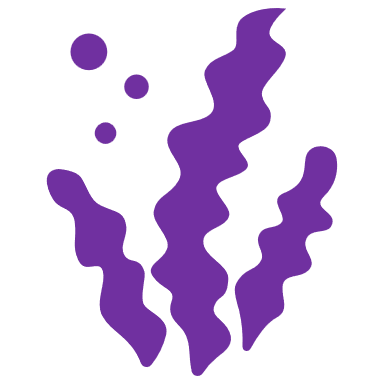 |
| *Nereocystis luetkeana* | 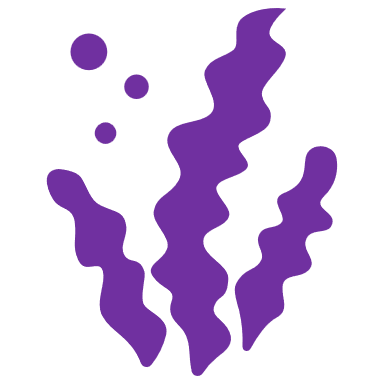 | 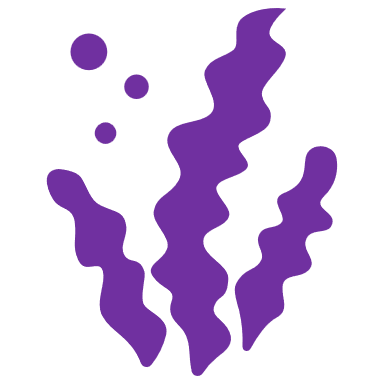 |
| *Pleurophycus gardneri* | 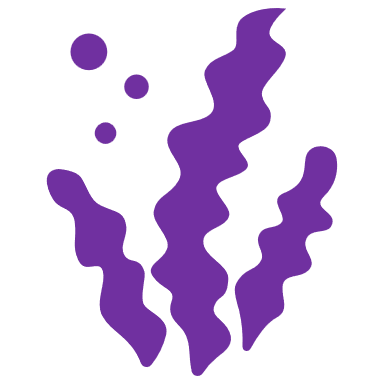 | 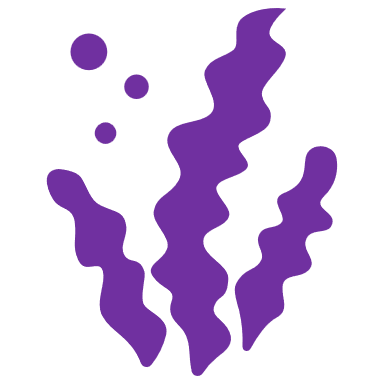 |
| *Pterygophora californica* | 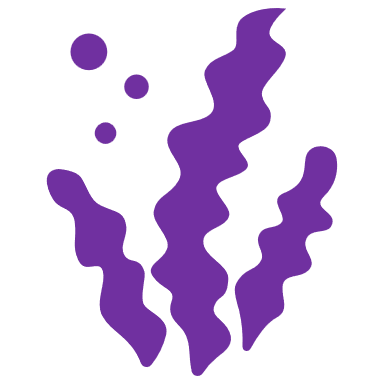 | 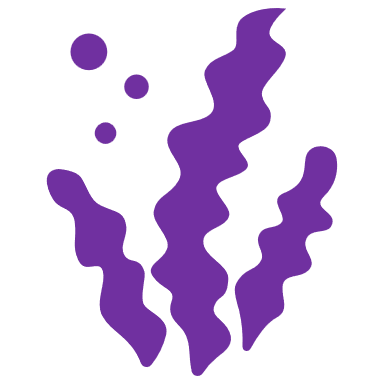 |
| *Saccharina latissima* | 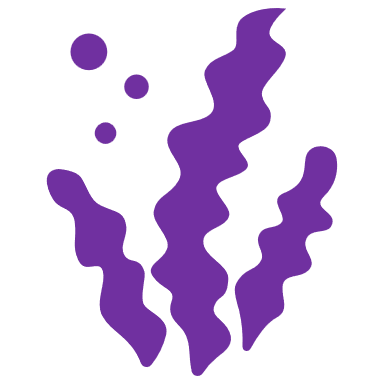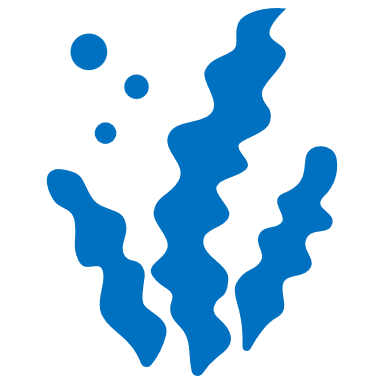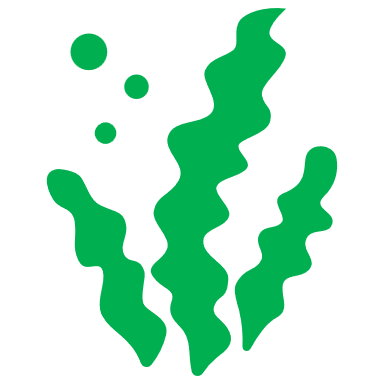 | 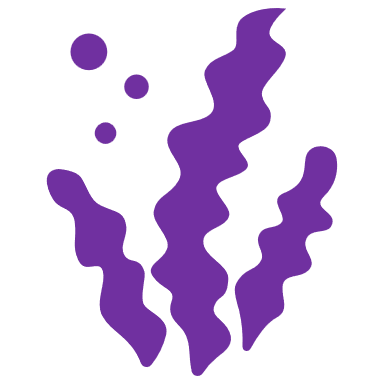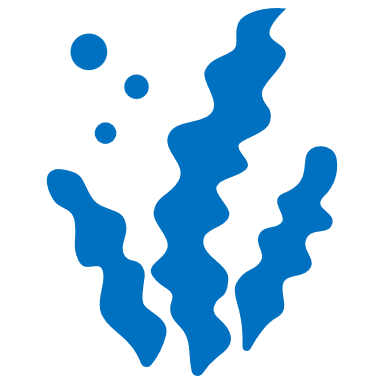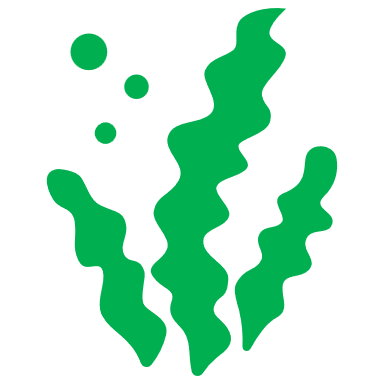 |

**Table C3:** Potential extent (Mha) of kelp forests in Canada. All estimates are presented as the potential lower bound (25^th^ percentile), median, upper bound (75% percentile), and maximum potential areas of kelp forests on each coast.

| **Ocean** | **Extent Estimates (Mha)** | | **Source** |
| --- | --- | --- | --- |
| **Pacific** |  |  |  |
| *Surface*  *kelp forests* | Maximum | 0.29 | Rocky reef above 10m |
|  | Upper Bound | 0.11 | Rocky reef above 10 m intersecting historic surface canopy maps compiled from the BC Shore Zone oblique shoreline aerial survey (Howes et al., 1994) |
|  | Lower Bound | 0.005 | Recent surface canopy maps determined by detections at peak biomass from Sentinel-2 satellite imagery (Mora-Soto et al., 2020) |
| *Subsurface kelp forests* | Maximum | 0.49 | Rocky reef above 20 m (maximum kelp cover) |
|  | Upper Bound | 0.19 | Rocky reef above 20m (75^th^ percentile kelp cover) |
|  | Median | 0.09 | Rocky reef above 20m (median kelp cover) |
|  | Lower Bound | 0.02 | Rocky reef above 20m (25^th^ percentile kelp cover) |
| **Arctic** |  |  |  |
| *Subsurface kelp forests* | Maximum | 4.55 | Rocky reef above 20m (maximum kelp cover) |
|  | Upper Bound | 3.2 | Rocky reef above 20m (75^th^ percentile kelp cover) |
|  | Median | 1.64 | Rocky reef above 20m (median kelp cover) |
|  | Lower Bound | 0.75 | Rocky reef above 20m (25^th^ percentile kelp cover) |
| **Atlantic** |  |  |  |
| *Subsurface kelp forests* | Maximum | 1.31 | Rocky reef above 20m (maximum kelp cover) |
|  | Upper Bound | 0.52 | Rocky reef above 20m (75^th^ percentile kelp cover) |
|  | Median | 0.14 | Rocky reef above 20m (median of kelp cover) |
|  | Lower Bound | 0.02 | Rocky reef above 20m (25^th^ percentile of kelp cover) |
| **Canada** |  |  |  |
| *Subsurface*  *kelp forests* | Maximum | 6.43 | Rocky reef above 20m (maximum kelp cover) |
|  | Upper Bound | 3.90 | Rocky reef above 20m (75^th^ percentile kelp cover) |
|  | Median | 1.86 | Rocky reef above 20m (median of kelp cover) |
|  | Lower Bound | 0.70 | Rocky reef above 20m (25^th^ percentile of kelp cover) |

**Table C4.** Pairwise conditional support for significant differences between the posterior mean estimated a) carbon stocks (Mg C Ha^-1^) and b) carbon production rates (Mg C ha^-1^ yr^-1^) of kelp species in Canada. Cells show the percent probability support that row species are significantly different from column species. Species names are abbreviated as followed: *Agarum clathratum / Neoagarum fimbriatum* (Ac / Nf), *Costaria costata* (Cc), *Laminaria digitata / Hedophyllum nigripes* (Ld / Hn), *Laminaria solidungula* (Ls), *Macrocystis pyrifera* (Mp), *Nereocystis luetkeana* (Nl), *Pterygophora californica* (Pc), *Pleurophycus gardneri* (Pg), and *Saccharina latissima* (Sl).

| **a) Carbon Stocks**  **(Mg Ha^-1^)** | **Pacific** | | | | | | | | **Arctic** | | | | **Atlantic** | | |
| --- | --- | --- | --- | --- | --- | --- | --- | --- | --- | --- | --- | --- | --- | --- | --- |
|  | Ac / Nf | Cc | Ld / Hn | Mp | Nl | Pc | Pg | Sl | Ac / Nf | Ld / Hn | Ls | Sl | Ac / Nf | Ld / Hn | Sl |
| **Pacific** | | | | | | | | | | | | | | | |
| Ac / Nf | NA | 52.2 | 18.1 | 0.0 | 2.7 | 0.1 | 61.5 | 21.6 | 0.0 | 1.9 | 6.1 | 0.1 | 47.5 | 26.0 | 22.6 |
| Cc | 47.8 | NA | 17.4 | 0.0 | 2.7 | 0.1 | 64.1 | 21.1 | 0.0 | 1.8 | 2.5 | 0.1 | 46.7 | 25.6 | 22.4 |
| Ld / Hn | 81.9 | 82.6 | NA | 1.8 | 6.0 | 11.3 | 83.2 | 49.4 | 33.1 | 16.3 | 65.1 | 1.7 | 76.2 | 48.1 | 31.5 |
| Mp | 100.0 | 100.0 | 98.2 | NA | 28.5 | 89.9 | 100.0 | 98.3 | 100.0 | 82.2 | 100.0 | 22.1 | 100.0 | 95.2 | 60.1 |
| Nl | 97.3 | 97.3 | 94.0 | 71.4 | NA | 86.2 | 97.4 | 93.8 | 93.9 | 83.4 | 96.2 | 50.0 | 96.9 | 92.3 | 68.7 |
| Pc | 99.9 | 99.9 | 88.7 | 10.2 | 13.8 | NA | 99.9 | 86.3 | 91.9 | 51.6 | 99.1 | 6.9 | 98.9 | 81.3 | 47.0 |
| Pg | 38.6 | 35.8 | 16.8 | 0.0 | 2.6 | 0.1 | NA | 20.2 | 0.1 | 1.8 | 6.3 | 0.1 | 43.5 | 24.9 | 22.1 |
| Sl | 78.4 | 79.0 | 50.6 | 1.7 | 6.2 | 13.7 | 79.7 | NA | 36.8 | 18.2 | 63.6 | 1.7 | 74.1 | 48.9 | 31.7 |
| **Arctic** | | | | | | | | | | | | | | | |
| Ac / Nf | 100.0 | 100.0 | 66.9 | 0.0 | 6.1 | 8.1 | 99.9 | 63.2 | NA | 17.2 | 96.9 | 0.9 | 95.5 | 58.2 | 34.9 |
| Ld / Hn | 98.1 | 98.2 | 83.7 | 17.7 | 16.6 | 48.4 | 98.2 | 81.8 | 82.8 | NA | 94.9 | 10.8 | 96.1 | 77.7 | 47.2 |
| Ls | 93.9 | 97.5 | 34.9 | 0.0 | 3.8 | 0.9 | 93.7 | 36.4 | 3.1 | 5.1 | NA | 0.2 | 74.1 | 37.0 | 27.0 |
| Sl | 99.9 | 99.9 | 98.3 | 77.9 | 50.0 | 93.1 | 99.9 | 98.3 | 99.1 | 89.2 | 99.8 | NA | 99.8 | 96.6 | 70.2 |
| **Atlantic** | | | | | | | | | | | | | | | |
| Ac / Nf | 52.5 | 53.3 | 23.8 | 0.0 | 3.1 | 1.1 | 56.5 | 25.9 | 4.5 | 3.9 | 25.9 | 0.2 | NA | 29.0 | 23.4 |
| Ld / Hn | 74.0 | 74.4 | 51.9 | 4.8 | 7.7 | 18.7 | 75.1 | 51.1 | 41.8 | 22.3 | 63.1 | 3.4 | 71.0 | NA | 32.5 |
| Sl | 77.4 | 77.6 | 68.5 | 39.9 | 31.3 | 53.1 | 77.9 | 68.2 | 65.1 | 52.8 | 73.0 | 29.8 | 76.6 | 67.5 | NA |
| **b) Carbon Production (Mg Ha^-1^ Yr^-1^)** | **Pacific** | | | | | | | | **Arctic** | | | | **Atlantic** | | |
|  | Ac / Nf | Cc | Ld / Hn | Mp | Nc | Pc | Pg | Sl | Ac / Nf | Ld / Hn | Ls | Sl | Ac / Nf | Ld / Hn | Sl |
| **Pacific** | | | | | | | | | | | | | | |  |
| Ac / Nf | NA | 52.4 | 10.9 | 0.3 | 7.0 | 31.1 | 14.3 | 3.2 | 28.8 | NA | 53.8 | 24.3 | 47.1 | 29.8 | 37.0 |
| Cc | 47.6 | NA | 9.4 | 0.1 | 6.7 | 30.3 | 8.6 | 1.7 | 24.5 | NA | 77.0 | 23.0 | 43.9 | 29.6 | 37.0 |
| Ld / Hn | 89.1 | 90.6 | NA | 8.3 | 22.3 | 61.8 | 75.0 | 39.1 | 77.7 | NA | 90.8 | 67.7 | 86.0 | 48.3 | 54.7 |
| Mp | 99.7 | 99.9 | 91.6 | NA | 57.0 | 87.9 | 98.8 | 84.7 | 98.1 | NA | 99.9 | 96.5 | 99.1 | 72.4 | 76.0 |
| Nc | 93.0 | 93.3 | 77.7 | 43.0 | NA | 79.0 | 88.2 | 70.8 | 88.8 | NA | 93.3 | 85.2 | 91.9 | 65.9 | 70.2 |
| Pc | 68.9 | 69.6 | 38.1 | 12.0 | 21.0 | NA | 55.7 | 30.3 | 59.1 | NA | 69.8 | 51.3 | 67.0 | 41.5 | 46.8 |
| Pg | 85.7 | 91.4 | 25.0 | 1.2 | 11.8 | 44.3 | NA | 13.3 | 61.9 | NA | 91.6 | 47.1 | 78.4 | 37.6 | 45.5 |
| Sl | 96.8 | 98.3 | 60.9 | 15.3 | 29.2 | 69.7 | 86.7 | NA | 87.5 | NA | 98.3 | 77.7 | 93.7 | 54.2 | 60.3 |
| **Arctic** | | | | | | | | | | | | | | | |
| Ac / Nf | 71.2 | 75.5 | 22.2 | 1.9 | 11.2 | 40.9 | 38.1 | 12.5 | NA | NA | 76.0 | 40.0 | 66.1 | 35.6 | 42.9 |
| Ld / Hn | NA | NA | NA | NA | NA | NA | NA | NA | NA | NA | NA | NA | NA | NA | NA |
| Ls | 46.2 | 23.0 | 9.2 | 0.1 | 6.6 | 30.2 | 8.4 | 1.6 | 24.0 | NA | NA | 22.6 | 43.1 | 29.5 | 36.9 |
| Sl | 75.7 | 77.0 | 32.3 | 3.5 | 14.8 | 48.8 | 52.9 | 22.3 | 60.0 | NA | 77.3 | NA | 72.4 | 40.0 | 47.0 |
| **Atlantic** | | | | | | | | | | | | | | | |
| Ac / Nf | 53.0 | 56.1 | 14.0 | 0.9 | 8.1 | 33.0 | 21.6 | 6.3 | 33.9 | NA | 56.9 | 27.7 | NA | 30.6 | 37.3 |
| Ld / Hn | 70.2 | 70.4 | 51.7 | 27.6 | 34.1 | 58.4 | 62.4 | 45.8 | 64.4 | NA | 70.6 | 60.0 | 69.4 | NA | 54.1 |
| Sl | 63.1 | 63.0 | 45.3 | 24.0 | 29.8 | 53.3 | 54.5 | 39.7 | 57.1 | NA | 63.1 | 53.0 | 62.7 | 45.9 | NA |

**Table C5**. Model definitions for carbon standing stocks and annual accumulation rates associated with kelp forest found. All models include site and source ID as random effects.

| **Model** | **Model definition** |
| --- | --- |
| *Carbon standing stocks (CS)* | |
| CS.1 | carbon stock ~ 1 |
| CS.2 | carbon stock ~ ocean temperature |
| CS.3 | carbon stock ~ ocean^*^ |
| CS.4 | carbon stock ~ ocean temperature + ocean^*^ |
| *Carbon accumulation rates (CP)* | |
| CP.1 | carbon accumulation ~ 1 |
| CP.2 | carbon accumulation ~ ocean temperature |
| CP.3 | carbon accumulation ~ ocean^*^ |
| CP.4 | carbon accumulation ~ ocean temperature + ocean^*^ |

**^*^***Included only for species that occur across multiple oceans.*

**Table C6**. Prior definitions for all carbon stock and carbon accumulation rate models.

| **Coefficient** | **Prior** | **Notes** |
| --- | --- | --- |
| Intercept | Student_t(3,0,9) | Scaled to the range of observed carbon stocks and accumulation rates found in Canadian kelp species |
| Slope | Student_t(3,0,9) | For fixed effects |
| Sigma | Student_t(3,0,9) | For random effects |

**Table C7.** Results of leave-one-out pareto-smoothed importance sampling (LOO-PSIS) model evaluation for carbon stocks of kelp forest-forming species. All models include site and source identity as random effects. Elpd_diff is the difference in the expected log predictive density between model structures; se_diff is the difference in standard error explained between models. Guidelines for interpreting these criteria are available at <https://mc-stan.org/loo/reference/loo-glossary.html>.

| **a) Carbon Standing Stocks (Mg Ha-1)** | | | | |
| --- | --- | --- | --- | --- |
| **Taxa** | **Model** | **Effects tested** | **elpd_diffo** | **se_diff** |
| *Macrocystis pyrifera* | CS.1 | carbon stock ~ 1 | 0 | 0 |
|  | CS.2 | carbon stock ~ ocean temperature | -0.1 | 0.5 |
| *Nereocystis leutkeana* | CS.2 | carbon stock ~ ocean temperature | 0 | 0 |
|  | CS.1 | carbon stock ~ 1 | -0.7 | 0.4 |
| *A. clathratum / N. fimbriatum* | CS.2 | carbon stock ~ ocean temperature | 0 | 0 |
|  | CS.3 | carbon stock ~ ocean | -0.8 | 1.8 |
|  | CS.1 | carbon stock ~ 1 | -1.5 | 1.7 |
|  | CS.4 | carbon stock ~ ocean temperature + ocean | -2.3 | 1.17 |
| *Costaria costata* | CS.1 | carbon stock ~ 1 | 0 | 0 |
|  | CS.2 | carbon stock ~ ocean temperature | -0.9 | 0.6 |
| *L. digitata / H. nigripes* | CS.2 | carbon stock ~ ocean temperature | 0 | 0 |
|  | CS.3 | carbon stock ~ ocean | -0.4 | 0.7 |
|  | CS.4 | carbon stock ~ ocean temperature + ocean | -0.4 | 0.7 |
|  | CS.1 | carbon stock ~ 1 | -0.7 | 1.1 |
| *Laminaria solidungula* | CS.1 | carbon stock ~ 1 | 0 | 0 |
|  | CS.2 | carbon stock ~ ocean temperature | -0.9 | 0.2 |
| *Pleurophycus gardneri* | CS.1 | carbon stock ~ 1 | 0 | 0 |
|  | CS.2 | carbon stock ~ ocean temperature | -0.8 | 0.8 |
| *Pterygophora californica* | CS.1 | carbon stock ~ 1 | 0 | 0 |
|  | CS.2 | carbon stock ~ ocean temperature | -0.3 | 1.2 |
| *Saccharina latissima* | CS.4 | carbon stock ~ ocean temperature + ocean | 0 | 0 |
|  | CS.1 | carbon stock ~ 1 | -1.2 | 1 |
|  | CS.3 | carbon stock ~ ocean | -3.6 | 1.7 |
|  | CS.2 | carbon stock ~ ocean temperature | -3.7 | 2.1 |
| **b) Carbon Accumulation Rates (Mg Ha-1 Yr-1)** | | | | |
| **Taxa** | **Model** | **Effects tested** | **elpd_diff** | **se_diff** |
| *Macrocystis pyrifera* | CP.1 | carbon accumulation ~ 1 | 0 | 0 |
|  | CP.2 | carbon accumulation ~ ocean temperature | -0.5 | 0.3 |
| *Nereocystis leutkeana* | CP.2 | carbon accumulation ~ ocean temperature | 0 | 0 |
|  | CP.1 | carbon accumulation ~ 1 | -0.1 | 0.4 |
| *A. clathratum / N. fimbriatum* | CP.1 | carbon accumulation ~ 1 | 0 | 0 |
|  | CP.2 | carbon accumulation ~ ocean temperature | -0.2 | 0.6 |
|  | CP.3 | carbon accumulation ~ ocean | -0.6 | 0.4 |
|  | CP.4 | carbon accumulation ~ ocean temperature + ocean | -0.7 | 0.5 |
| *Costaria costata* | CP.1 | carbon accumulation ~ 1 | 0 | 0 |
|  | CP.2 | carbon accumulation ~ ocean temperature | -0.9 | 0.4 |
| *L. digitata / H. nigripes* | CP.2 | carbon accumulation ~ ocean temperature | 0 | 0 |
|  | CP.1 | carbon accumulation ~ 1 | -0.1 | 0.6 |
|  | CP.3 | carbon accumulation ~ ocean | -0.5 | 0.6 |
|  | CP.4 | carbon accumulation ~ ocean temperature + ocean | -0.6 | 0.4 |
| *Laminaria solidungula* | CP.1 | carbon accumulation ~ 1 | 0 | 0 |
|  | CP.2 | carbon accumulation ~ ocean temperature | 1.2 | 0.4 |
| *Pleurophycus gardneri* | CP.1 | carbon accumulation ~ 1 | 0 | 0 |
|  | CP.2 | carbon accumulation ~ ocean temperature | -1.2 | 0.4 |
| *Pterygophora californica* | CP.1 | carbon accumulation ~ 1 | 0 | 0 |
| *Saccharina latissima* | CP.4 | carbon accumulation ~ ocean temperature + ocean | 0 | 0 |
|  | CP.2 | carbon accumulation ~ ocean temperature | -2.1 | 2.5 |
|  | CP.3 | carbon accumulation ~ ocean | -3.6 | 2.3 |
|  | CP.1 | carbon accumulation ~ 1 | -3.8 | 2.4 |
